# Divergent strategies to reduce stomatal pore index during water deficit in perennial angiosperms

**DOI:** 10.1101/2020.07.07.191817

**Authors:** Noel Anthony Mano, Santiago Franco Lopez, Michael V. Mickelbart

**Author notes:** Author for correspondence: *Michael V. Mickelbart, Email:.

## Abstract

⍰ Modulation of stomatal development may be an acclimation response to low water availability. However, stomatal development plasticity has been assessed in very few species.

⍰ We quantified leaf anatomy traits, including stomatal index (SI), density (SD), size (SS), and pore index (SPI), in response to water-deficit stress in river birch (*Betula nigra* L.), eastern redbud (*Cercis canadensis* L.), and silver maple (*Acer saccharinum* L.).

⍰ Birch and redbud, but not maple, had reduced SPI in response to water deficit. The mechanism by which SPI reduction occurred (via SD or SS) varied among species and with severity of water stress. Despite reduced SPI in birch and redbud, anatomical changes were relatively small and had a minor to no effect on the theoretical maximum stomatal conductance. Furthermore, gas-exchange rates were equivalent to well-watered plants following media re-irrigation.

⍰ In some tree species, stomatal development is downregulated in response to water deficit conditions. Stomatal development plasticity is facilitated by smaller or fewer stomata, depending upon the species and the intensity of the stress. Water-deficit-induced plasticity in stomatal development is species-specific, likely due to species adaptation to ecological niches.

## Introduction

Understanding plant acclimation and adaptation to water deficit will become increasingly important as climate change increases the frequency and intensity of drought. Drought events are projected to increase in frequency globally, and in the US Midwest in particular, from once every five years to once every other year by 2050 (Jin *et al.*, 2018). In natural ecosystems, drought events result in reduced ecosystem productivity (Wu *et al.*, 2011) and plant mortality (Gitlin *et al.*, 2006; Klos *et al.*, 2009). Reduced water availability constrains CO_2_ assimilation and reduces carbon provisioning from shoots to roots (Ruehr *et al.*, 2009). Drought events may decrease tree water potentials to below the zero-carbon assimilation point (Breshears *et al.*, 2009), which can deplete tree carbon reserves and result in tree death (McDowell, 2011). Although there is evidence for large-scale ecosystem resilience to dynamic seasonal availability of water (Ponce-Campos *et al*., 2013), the role of leaf hydrological plasticity in achieving these acclimations is unclear.

The capacity of stomata to regulate water loss during periods of water deficit is critical to plant growth and survival. Stomatal closure is an important transitory response during drought events, but during chronic water deficit, modulation of stomatal anatomy may also be important for water conservation (Liu *et al.*, 2018). Smaller stomata close more quickly in response to decreased leaf hydration status than larger stomata (Drake *et al.*, 2013; Giday *et al.*, 2013) and smaller guard cells are biomechanically optimized to open at lower turgor pressure (Spence *et al.*, 1986) but without increased carbon costs (Raven, 2014) enabling maintenance of CO_2_ assimilation during water-stress events.

In dicotyledonous plants, stomatal development primarily occurs during early leaf emergence, when developing leaves are composed of dividing and differentiating stem cells (Rawson & Craven, 1975; Andriankaja *et al.*, 2012), although the specific timing of development varies among species (Woodall *et al.*, 1998). Stomatal differentiation can occur late in leaf development in some species (Ludlow, 1991), and there is some evidence for the persistence of meristemoids that can form new stomata late in leaf development (Geisler *et al.*, 2000). Within the first two days of leaf emergence, leaf veins form (Kang & Dengler, 2004) and vein density is subsequently modulated by passive dilution during leaf expansion (Carins Murphy *et al.*, 2012). Based on this developmental timing, environmental effects on the ultimate anatomy of a leaf are likely to occur early in the development of that leaf. A stress occurring during the expansionary phase of leaf development may result in changes in stomatal density due to altered cell turgor and therefore cell size, but the relative numbers of guard cells to pavement cells is unlikely to be significantly altered because cell identity is mostly fixed by this stage. This illustrates the difficulty in drawing conclusions on mechanistic responses to diverse environments from ecological experiments, such as those involving stands of trees that receive different precipitation levels (Reyer *et al.*, 2013), since it is difficult to ascertain when a water deficit event occurs relative to leaf development.

Stomatal development is sensitive to environmental cues at the cellular level, particularly light (Casson *et al.*, 2009; Kang *et al.*, 2009) and CO_2_ (Woodward, 1987; McElwain & Chaloner, 1995; Franks & Beerling, 2009), suggesting that the modulation of stomatal development is an acclimation trait. The expression of several genes that regulate stomatal development are downregulated in response to osmotic stress (Kilian *et al.*, 2007; Harb *et al.*, 2010; Yoo *et al.*, 2010; Baerenfaller *et al.*, 2012; Kumari *et al.*, 2014; Yoo *et al.*, 2019), and have been linked to a concomitant suppression of stomatal development in *Arabidopsis* (Skirycz *et al.*, 2011; Kumari *et al.*, 2014; Yoo *et al.*, 2019) and soybean (Tripathi *et al.*, 2016). Similar gene expression patterns may exist in tree species, but very few data have been collected (Hamanishi *et al.*, 2012; Viger *et al.*, 2016).

Despite these observed links between water deficit and stomatal plasticity at the anatomical and molecular levels, plasticity of SI, stomatal density (SD), and stomatal size (SS) in response to water deficit varies widely among (de Silva *et al.*, 2012; Hamanishi *et al.*, 2012) and within (Pääkkönen *et al.*, 1998; Hovenden & Vander Schoor, 2012) tree species. Between-species variation may be explained by different degrees and methods of water deficit imposed by different authors, such as withholding water over a period of time (Hamanishi *et al.*, 2012) or different degrees of maintained media water content (de Silva *et al.*, 2012; Aasamaa *et al.*, 2001; Catoni *et al.*, 2017). Within-species differences in stomatal development plasticity could be caused by different provenances having different stomatal patterning as an adaptation to local levels of moisture (Dunlap & Stettler, 2001; Pearce *et al.*, 2006, McKown *et al.*, 2014).

Stomatal pore index (SPI, Sack *et al.*, 2003) and theoretical maximum stomatal conductance (*g_smax_*, Franks & Farquhar, 2001) are derived traits that combine the frequency and dimensions of stomata to describe the effective potential water loss from the leaf interior, which effectively predicts leaf gas exchange (Dow *et al.*, 2014; McElwain *et al.*, 2016). These traits account for all the possible changes in stomatal anatomy that could occur in producing leaves optimized for water conservation. For example, simply reporting a lack of SD plasticity may omit the production of smaller stomata that leads to reduced SPI and/or *g_smax_*. Since SPI and *g_smax_* are frequently not reported, conclusions cannot yet be drawn about the role of stomatal development plasticity in facilitating water deficit tolerance in trees.

We hypothesized that perennial species exposed to persistent water-deficit stress acclimate via a reduction in stomatal development and overall stomatal coverage to minimize water loss. We examined three tree species: *Betula nigra* L., *Acer saccharinum* L., and *Cercis canadensis* L., chosen to represent the Betulaceae, Sapindaceae, and Fabaceae families, respectively. These species occupy a large range of urban, rural, and forested land across temperate North America.

## Materials and methods

### Plant materials and growth conditions

One-year-old bare-root river birch *(Betula nigra* L.) were planted in 8.5 L containers, and four-week-old redbud *(Cercis canadensis* L.) and silver maple *(Acer saccharinum* L.) seedlings were planted in 3.4 L containers in a BM8 Berger soilless substrate (Berger, Saint-Modeste, QC, Canada). Plants were maintained in a greenhouse in the Purdue Horticulture Plant Growth Facility from June to December 2018. A minimum 14 h photoperiod was provided with 100-watt high-pressure sodium lamps. Temperature and relative humidity were measured using two HOBO^®^ data loggers (Onset Computer Corporation, Bourne, MA, USA), and the daily light integral was quantified with an external weather station (Fig. S1). Average day and night temperatures were 24.7 and 23.6°C, respectively, and average relative humidity (RH) was 70%. Vapor pressure deficit (VPD, kPa) was calculated as

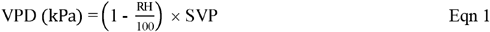

where SVP is the saturated vapor pressure at a given daily temperature, derived from standard tables of the two quantities.

Two experiments were performed, the first imposing a mild stress, and the second imposing a more severe stress. Media water content (MWC) was maintained by weighing plants and replacing water on a regular basis to 100 or 60% MWC (experiment 1), and 100 or 40% MWC (experiment 2) of the initial saturated weight (including initial plant biomass) and was calculated as

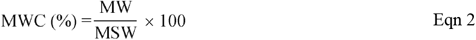

where MW is the weight of the container system (consisting of container, media, and plant) on a given day, and MSW is the saturated weight of the container system at the beginning of the experiment. For the first experiment, plants were irrigated to media capacity until establishment. After 95 d post-planting, water was withheld for 18 d until the target MWC of 60% was reached in the water-stressed (WS) treatment plants. Control well-watered (WW) plants were irrigated every 3–4 d as necessary, and WS plants every 24 h to 60% of their initial saturated weight (Fig. S2). In the second experiment, plants were irrigated to media capacity until establishment and after 54 d, WS was initiated. WW plants were irrigated every 2 d, and WS plants were irrigated every day (redbuds) or 2 d (maples) to 40% of their initial saturated weight (Fig. S2). Birch was not included in the 40% MWC treatment.

Acidified water was supplemented with water-soluble fertilizer (ICL Specialty Fertilizers, Dublin, OH, USA) to provide the following (in mg L^-1^): 150 N, 9.8 P, 119 K, 12 Mg, 21 S, 1.5 Fe, 0.4 Mn and Zn, 0.2 Cu and B, and 0.1 Mo. Nitrate and ammonium sources of nitrogen were provided as 61 and 39% total N, respectively. Irrigation water was supplemented with 93% sulfuric acid (Brenntag, Reading, PA, USA) at 0.08 mL L^-1^ to reduce alkalinity to 100 mg L^-1^ and pH to a range of 5.8 to 6.2.

### Measurements

Leaves that emerged after experimental days 20 and 34 in experiments 1 and 2, respectively, following the establishment of treatment MWC levels, were used for data collection (Fig. S3; Table S1). To quantify leaf development in experiment 1, leaf 2 was photographed next to a ruler with a digital camera *c*. every two days over 40 d and leaf area (LA) was quantified using ImageJ software (National Institutes of Health, Bethesda, MD, USA). A logistic curve was fit to the leaf area A at time *t* for each leaf (Fig. S4a-c):

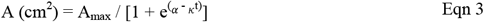

where α is the sigmoidal midpoint and *κ* is the logistic growth rate. The logistic curve was linearized and used to calculate the maximum leaf growth rate, experimental day on which the leaf reached 50% full expansion, and the number of days to full leaf expansion (Fig. S4d-f).

To confirm that leaves used for data collection were fully expanded, LA was assessed in leaves 2 (Fig. S4) and 4 in experiment 1. In experiment 2, the size of one leaf that had developed after the initiation of water deficit was measured over 7 days prior to harvest to ensure full expansion of this leaf. Final LA was quantified during the final destructive harvest, by removing the leaf and imaging against a white background.

Net CO_2_ assimilation (*A*, μmol CO_2_ m^-2^ s^-1^), stomatal conductance (*g_s_*, mol H_2_Om^-2^ s^-1^) and transpiration (*E*, mmol H_2_O m^-2^s^-2^) were measured using a portable gas-exchange analyzer (LI-6400XT; LI-COR Biosciences, Lincoln, NE, USA). Chamber conditions during measurements were 1800 μmol m^-2^ s^-1^ PAR, air temperature range of 29.0–32.8°C, and average VPD was 2.5 ± 0.4 kPa.

After scanning the leaf used for gas exchange measurements, a segment of leaf, including a portion of the midrib and petiole, was collected, weighed to obtain the fresh weight (FW) and placed into a 50 mL conical tube with 25 mL of water. After the leaf had been immersed for *c.* 8 h, the leaf was blotted dry and weighed to obtain the turgid weight (TW). The leaf was then dried at 45°C to a constant weight to obtain the dry weight (DW). The relative water content (RWC) was calculated as

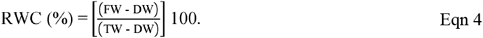

This portion of the leaf was also scanned for LA, from which the specific leaf weight was calculated as

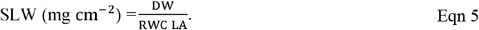

Osmotic potential was measured on the same leaves used for RWC. A leaf segment was removed, placed into an Eppendorf tube with a Costar Spin-X insert (Corning Incorporated, Corning, NY, USA), and immersed in liquid nitrogen. Samples were stored at −20°C. To extract cell sap, samples were thawed in sealed tubes for 5 min and then centrifuged for 5 min at 120 RPM x 100. Osmolality of a 10 μL of volume of cell sap was measured using a vapor pressure osmometer (VAPRO 5520; Wescor Inc., Logan, UT, USA). Osmolality was converted to osmotic potential (Ψ_*π*_) as

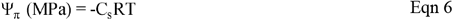

where C_s_ is osmolality, R is the gas constant, and T is temperature. The osmotic potential at full turgor (Ψ_*π*100_) was calculated as

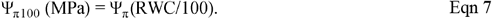

Epidermal traits were quantified from leaf impressions made on microscope slides using cyanoacrylate (Duro Super Glue; Henkel, Düsseldorf, Germany). Four images were taken from each impression with a DCM 900 microscope CMOS Camera (Oplenic Optronics, Hangzhou, China) of a 0.03 mm^2^ area, under 400X magnification using a light microscope (BH-2; Olympus, Tokyo, Japan). All three species are hypostomatous, so epidermal impressions were made only on the abaxial side of leaves. The number of stomata (SD) and pavement (PD) cells per unit area, and stomatal size (SS), equivalent to the area within the outer edges of the guard cell, was determined using ImageJ software. Only whole stomata and pavement cells bordering the top and right sides of each image were counted. Stomatal index (SI) was calculated as

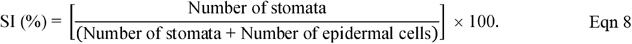

Pavement cell size was estimated by dividing the number of pavement cells counted in a region and dividing by the size of the visual field (30 000 μm^2^).

Stomatal size was calculated using the formula for an ellipse:

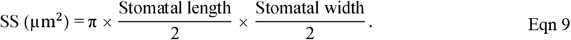

Stomatal pore index (SPI) was calculated as (Liu *et al*., 2018)

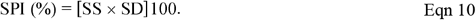

Theoretical maximum conductance (*g_smax_*) was calculated using the equation developed by Franks & Farquhar (2001)

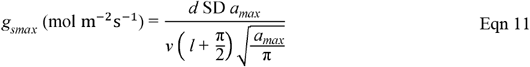

where *d* is the diffusivity of water in air (0.0000249 m^2^ s^-1^ at 25°C), *v* is the molar volume of air (0.0224 m^3^ mol^-1^), *l* is the stomatal pore depth equivalent to the width of a fully turgid guard cell, and *amax* is the maximum pore area, calculated using half the stomatal length as the pore length, as follows:

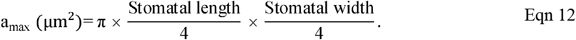

To quantify vein density (VD), the same leaf section from leaves 3–4 (experiment 1) and 1 (experiment 2) used for epidermal impressions was submerged in 50% sodium hypochlorite in a water bath at 50°C for 6 h. After the leaf was cleared, it was placed in a 0.0015% toluidine blue solution for 12 h, then observed under 40X magnification. Four images were taken per sample, each covering an area of 2.95 mm^2^ and avoiding the midvein of the leaf. Major and minor veins in the image were traced and the total length summed across all orders of veins. This sum was then divided by the area of the image to calculate VD as mm veins mm^-2^. Images were processed using ImageJ software.

### Statistical analysis

Experiments were conducted using a completely randomized design. In the first experiment, several measurements were made on multiple leaves from the same plant, and it was therefore analyzed as a repeated measures ANOVA. The leaf-treatment interaction was used to determine which leaves could be pooled for a given trait. Data was pooled across leaves if the interaction of treatment groups and leaves was not significant (*P* > 0.05). Regardless of pooling, each WS group was compared against the comparable control group by one-way ANOVA. Data were transformed with a Box-Cox transformation if needed to fulfill the assumption of normality. To calculate plasticity of anatomical traits, the natural logarithm of the response ratio was determined as ln(average of treatment group/average of control group), with the standard deviation of WW and WS groups combined to calculate the 95% confidence interval. A single leaf per plant was examined in the second experiment. All analyses were conducted in Jamovi v1.

## Results

### Plant growth and water status were adversely affected by water deficit

To verify the effects of the imposed WS, we quantified plant growth. Under WS, all trees were shorter than WW trees (Figs S5a, S6) and all trees produced smaller leaves except redbud, in which leaf size was not affected by 60% MWC (Figs S5b, S7). The 60% MWC treatment resulted in a reduced maximum leaf growth rate in all species (Fig. S4d), whereas the length of leaf development was longer only in maple leaves (Fig. S4e, f).

To assess treatment effects on plant water content, we quantified several water relations traits. Restricting MWC to 60% resulted in lower leaf RWC during the stress period, but this quickly recovered to WW levels following re-saturation of the media (Fig. S8). However, in the 40% MWC treatment, reduced RWC was observed in maple, but not redbud leaves (Fig. S8b, c). The 60% MWC WS level did not induce a reduction in osmotic potential (Ψ_π_) in any species, whereas Ψ_π_ was lower in maple and redbud leaves on trees grown under 40% MWC (Fig. S9a-c). In redbud leaves, this appears to be a passive effect, as Ψ_π100_ was not reduced (Fig. S9f).

### Water deficit resulted in plasticity of stomatal anatomy and leaf physiology

In these experiments, we quantified changes in leaf anatomy and physiology of three temperate tree species in response to WS. The plasticity of the various traits depended on species and the intensity of the water deficit treatment. Here we describe these changes for each of the five treatment combinations, relative to WW plants.

Under WS, birch trees produced leaves that were 45% smaller than WW leaves (Figs S5b, S7a), with similar leaf thickness (Fig. S10). Stomatal density was unchanged (Figs 1a, S11a) despite the production of 22% fewer stomata relative to the total cell population (Figs 1b, S12a). This is because stomata in WS leaves were 20% smaller (Figs 1c, S13a), primarily due to a decrease in stomatal length (as opposed to width) (Fig. S14a, d). Pavement cells were 26% smaller (Fig. S15a, b), so cell density of both types was unchanged (Fig. S16a, b), leading to the same stomatal distribution per unit area in all leaves. However, the smaller stomata in WS leaves resulted in a 17% lower SPI (Figs 1d, S17a). Despite the 20% reduction in SS, birch did not exhibit plasticity for *g_smax_* (Fig. 2a, b). There was no reduction in VD in WS birch leaves (Figs 3, S18a).

**Fig. 1.**
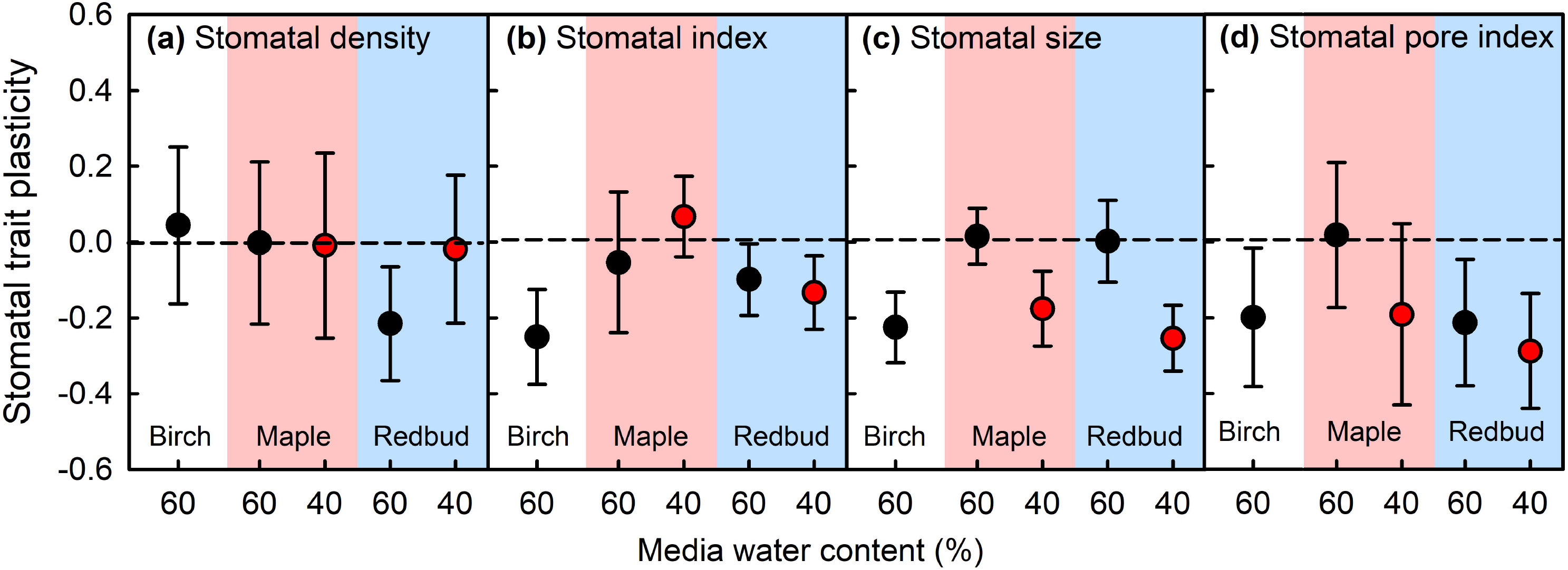
Plasticity of stomatal density (a), stomatal index (b), stomatal size (c), and stomatal pore index (d) following growth in containers with media water content (MWC) representing mild (60% MWC, black symbols) or severe (40%, red symbols) water stress (WS). Plasticity was calculated as the ln response ratio. Data is presented for the pooled second through fourth leaves that developed under 60% MWC (n = 4–6) and the first leaf that developed under 40% MWC (n = 6–8). Error bars represent 95% confidence intervals. WS plants are significantly different from well-watered plants if bars do not overlap 0.

**Fig. 2.**
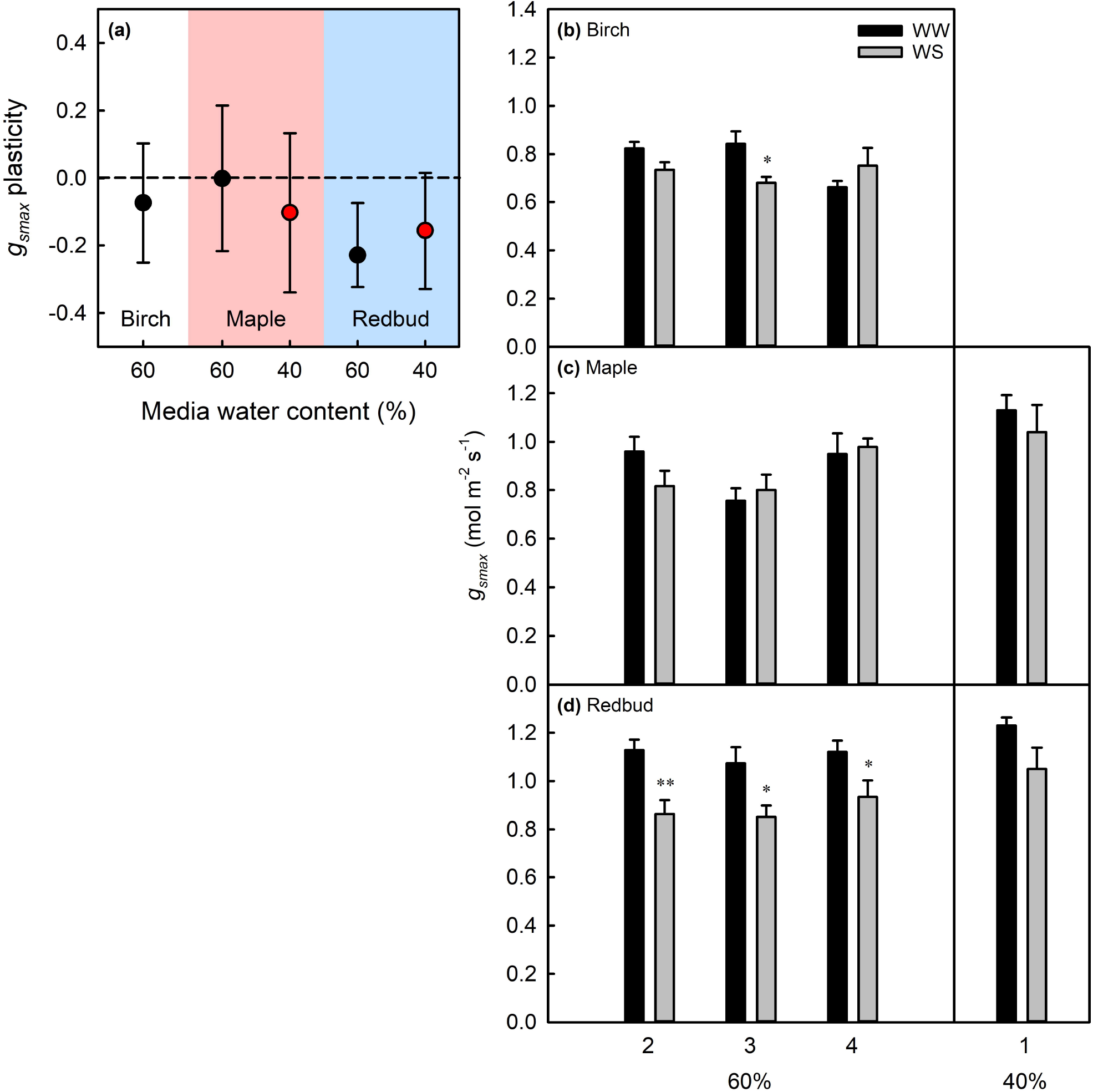
Plasticity of theoretical maximum stomatal conductance (a) following growth in containers with media water content (MWC) representing mild (60% MWC, black symbols) or severe (40%, red symbols) water stress (WS). Plasticity was calculated as the ln response ratio. Data is presented for the pooled third and fourth leaves that developed under 60% MWC (*n* = 4–6) and the first leaf that developed under 40% MWC (*n* = 6–8). Error bars represent 95% confidence intervals. WS plants are significantly different from well-watered plants if bars do not overlap 0. Data for individual leaves (b–d) are means ± SE, and * or ** denote a significant difference between well-watered (WW) and WS leaves at *P* < 0.05 or 0.01, respectively, based on one-way ANOVA (*n* = 4–6 for 60% MWC, *n* = 6–8 for 40% MWC).

Under WS conditions, *g_s_* was reduced by 90% in leaf 3 (in which *g_smax_* was reduced) and 76% in leaf 4 (in which *g_smax_* was unchanged) (Fig. 4a). During the water deficit stress, *A* was also 99 and 68% lower (Fig. 4d) and transpiration rate was reduced by 84 and 67% (Fig. S19a) in birch leaves 3 and 4, respectively. Due to stomatal closure, reduced SPI and smaller leaf size, WS birch leaves lost 85% less water during the water-deficit period (Fig. S19d). *g_s_* and *A* eventually recovered to levels comparable to WW leaves seven days after media re-saturation (Figs 4a, d). Despite the smaller LA and SS, whole-leaf transpiration also recovered to WW levels in this time frame (Fig. S19d). It is possible that such a recovery could occur because in leaf 4 SPI was not different in WW and WS leaves (Fig. S17a), and the size of WS leaf 4 was not as reduced as previously developed leaves (Fig. S7a).

In response to 60% MWC, maple leaves were 48% smaller (Figs S5b, S7b) and 18% thinner (Fig. S10) than WW leaves. However, epidermal anatomy was mostly non-plastic, with no change in SS, SI, or SD (Figs 1, S11b, S12b, S13b). There was also no change in pavement cell size (Fig. S15a, c) or density (Fig. S16a, c). As a result, neither SPI nor *g_smax_* were lower in WS leaves (Figs 1d, 2a, c, S17b). Maple leaf VD was also unchanged under 60% MWC (Figs 3, S18b).

**Fig. 3.**
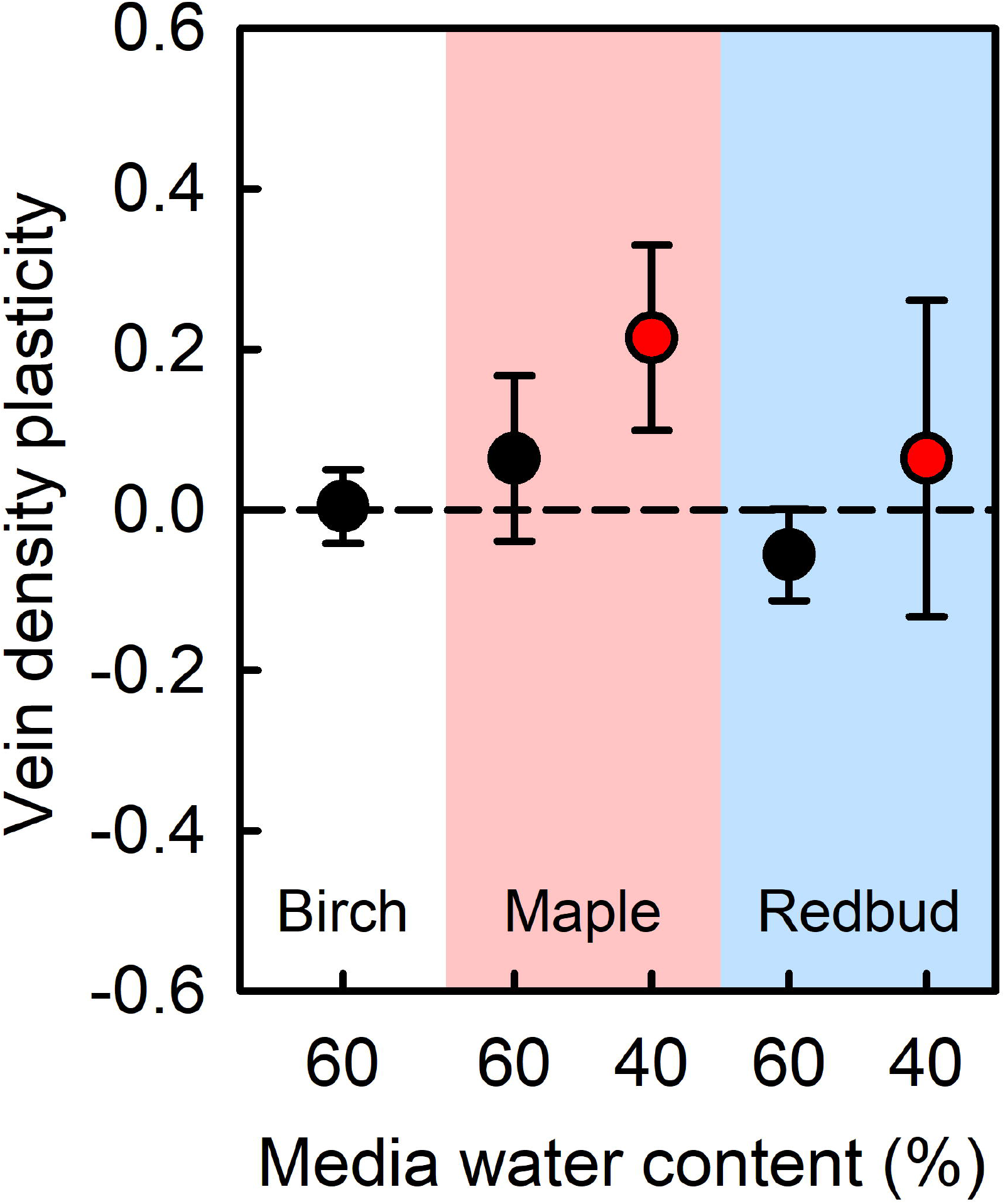
Plasticity of vein density following growth in containers with media water content (MWC) representing mild (60% MWC, black symbols) or severe (40%, red symbols) water stress (WS). Plasticity was calculated as the ln response ratio. Data is presented for the pooled third and fourth leaves that developed under 60% MWC (n = 4–6) and the first leaf that developed under 40% MWC (n = 6–8). Error bars represent 95% confidence intervals. WS plants are significantly different from well-watered plants if bars do not overlap 0.

Despite the lack of anatomical plasticity, *g_s_* and *A* were 83% lower overall in WS leaves, relative to WW leaves (Fig. 4b, e). Maple WS leaves 3 and 4 grown under 60% MWC had reduced *g_s_, A,* and *E* of 90 and 77% (Fig. 4b), 87 and 80% (Fig. 4e), and 78 and 72% (Fig. S19b), respectively. Due to stomatal closure and smaller leaf size, WS maple leaves lost 88% less water during the water-deficit period (Fig. S19e). *A* and *E* eventually recovered to levels comparable to WW leaves three days after media re-saturation (Figs 4e, S19b). Due to the smaller leaf size and stomatal closure, WS leaves lost 88% less water, relative to WW leaves, and whole-leaf transpiration was still depressed in WS leaves at the end of the recovery period (Fig. S19e).

**Fig. 4.**
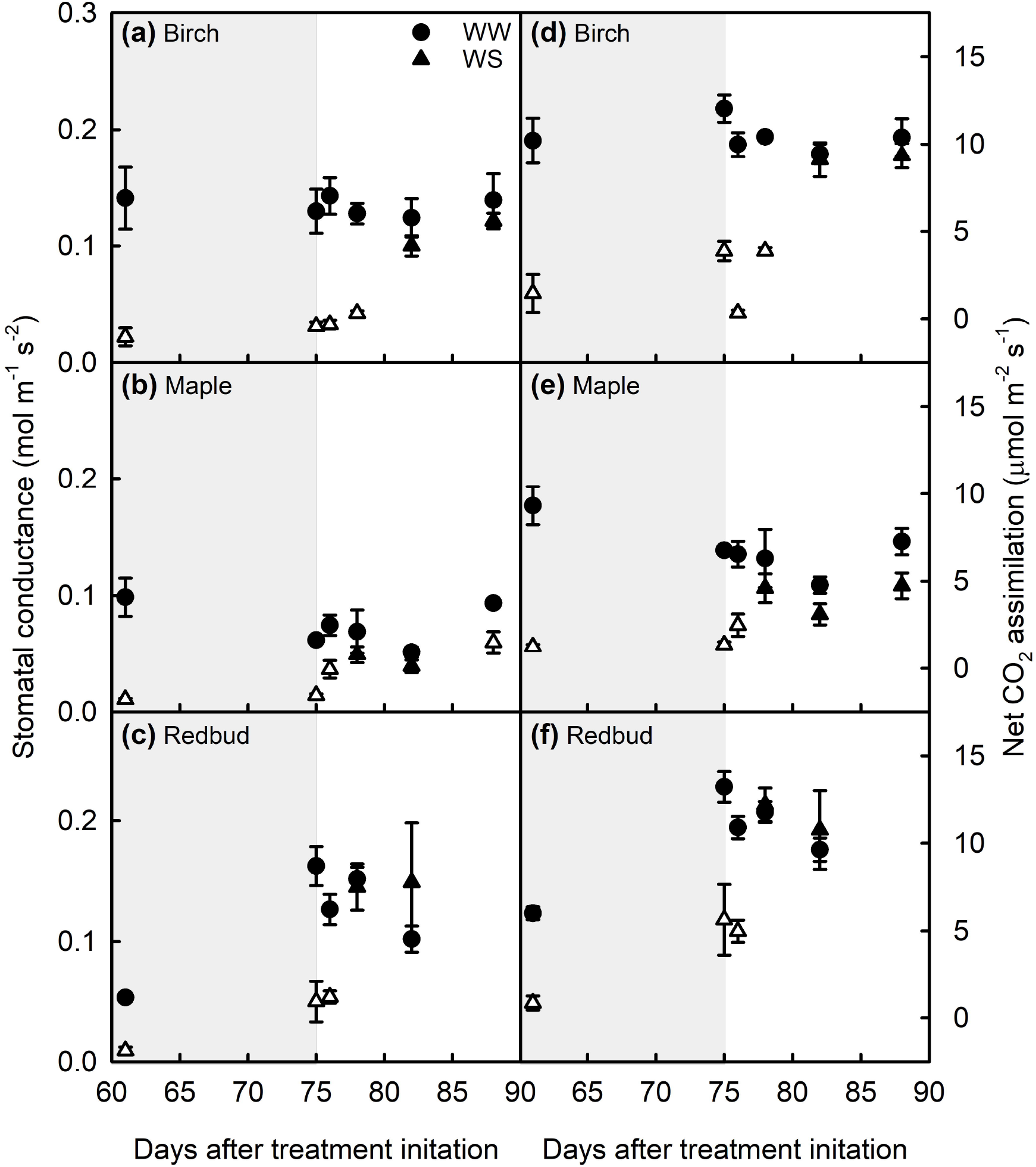
Stomatal conductance (a–c) and net CO_2_ assimilation (d–f) of leaves that developed under well-watered (WW, circles) or water-stress (WS, triangles) conditions (60% MWC) or conditions. Background shading represents maintained MWC treatments over time (light grey) and re-irrigation (white). Leaf 3 was used for day 61 measurements and leaf 4 for all other measurements (see Fig. S3). Data are means ± standard error (SE). Open symbols indicate a significant difference between WS and WW plants at *P* < 0.05, based on a one-factor ANOVA (*n* = 4–6).

Under the severe 40% MWC WS, maple leaves were 77% smaller (Figs S5, S7b) with no change in leaf thickness (Fig. S10). Due to a reduction in length (Fig. S14b), stomata in WS leaves were 13% smaller than those of WW leaves (Figs 1c, S13b). As in the 60% MWC treatment, no changes in stomatal anatomy occurred (Figs 1, 2, S11b, S12b, S17b). However, there was a 24% increase in VD (Figs 3, S18b). The 40% MWC WS resulted in a 50, 71, and 42% reduction in *g_s_, A,* and *E*, respectively (Fig. S20a-c). Overall, 40% MWC WS maple leaves lost 85% less water than WW leaves (Fig. S20d).

Redbud trees produced leaf types depending on the intensity of the WS treatment. Under the 60% MWC WS, redbud leaves were not smaller (Figs S5b, S7c), but they were 11% thinner (Fig. S10). There was no change in SS (Figs 1c, S13c, S14c, f), but SI was reduced by 10% (Figs 1b, S12c). Because epidermal cell size and density were unchanged (Figs S15a, d; S16a, d), this decrease in stomatal development resulted in a 20% decrease in SD (Figs 1a, S11c). With fewer stomata in redbud leaves, SPI and *g_smax_* were reduced by 20% (Figs 1d, 2a, d, 17c). Vein density was not different between WS and WW leaves (Figs 3, S18c).

Under the 60% MWC WS, *g_s_* was reduced by 87% in leaf 3 and 72% in leaf 4 (Fig. 4c), as both leaves had reduced *g_smax_. A* was also 86 and 61% lower during the water deficit period in these leaves (Fig. 4f). *E* was 85 and 67% lower (Fig. S19c). Although LA was unchanged, stomatal closure and lower SD resulted in WS redbud leaves losing 75% less water during the water-deficit period (Fig. S19f). Despite reduced SD and *g_smax_* in leaf 4, *g_s_, A, E,* and whole-leaf transpiration eventually recovered to levels comparable to WW redbud leaves three days after root zone re-saturation (Figs 4c, f; S19c, f).

By contrast, the severe (40% MWC) WS resulted in smaller redbud leaves (Figs S5b, S7c), but SLW was similar in WW and WS leaves (Fig. S10). In this case, redbud stomata were 23% smaller (Figs 1c, S13c), because of both shorter and narrower guard cell dimensions (Fig. S14c, f). Despite SI being reduced by 12% in these leaves (Figs 1b, S12c), SD was unchanged (Figs 1a, 11c). Because of smaller stomata, SPI was reduced by 25% (Figs 1d, S17c), but this was not sufficient to reduce *g_smax_* (Fig. S2a, d). However, *g_s_* and *A* were reduced (by 85 and 88%, respectively, Fig. S20a, b). Overall, the smaller LA and SS, as well as stomatal closure, meant that 40% MWC WS redbud leaves lost 93% less water than WW leaves (Fig. S20c, d).

## Discussion

### Effects of water stress on tree growth and development

Water-deficit treatments resulted in shorter trees in all species and smaller leaves in almost all species and treatments (Figs S5–S7). Similar to previous studies, birch LA was reduced with only a small effect on plant size under the mild stress we applied (Kleczewski *et al.*, 2012). In maple and redbud, the severe 40% MWC treatment resulted in a more dramatic reduction in growth compared to the 60% MWC treatment (Fig. S5).

Increased leaf thickness under water deficit conditions may increase the diffusion path from the leaf interior to the environment, thereby minimizing water loss from leaves (Syvertsen *et al.*, 1995; Sobrado, 2007). As noted in previous studies (Kleczewski *et al.*, 2012), there was no change in leaf thickness of river birch leaves in response to drought stress. Both increases and decreases in water-deficit-induced leaf thickness have been reported in related species *B. ermanii* (Kitao *et al.*, 2003; Tabata *et al.*, 2010) and *B. pendula* (Possen *et al.*, 2011; Aspelmeier & Leuschner, 2006). Genotypes of *Cercis canadensis* adapted to drier regions have thicker leaves than those adapted to wetter regions (Donselman & Flint, 1982; Abrams, 1988; Tipton & White, 1995; Fritsch *et al.*, 2018), but there are no reports of temporal water-deficit events on leaf thickness in redbud. In the present study, redbud leaves were thinner in response to the 60% MWC and were unchanged in response to the 40% MWC treatment, and the same pattern existed in maple. Decreased water availability also resulted in thinner leaves in related species *A. truncatum* (Li *et al.*, 2017) and *A. davidii* (Guo *et al.*, 2019). Our data show that the reduction in leaf thickness that occurs under the mild stress is not enhanced by the more severe water deficit. In redbud and maple, thinner leaves may be produced in response to mild water deficit to allow for easier hydration of the leaf. It is unclear why this response would not exist under more severe stress.

Decreasing water availability results in decreased RWC in tree species (Reddy *et al*., 2004) and experimental treatments that drastically reduce water availability can result in extremely low RWC (Ma *et al.*, 2015; Vieira *et al.*, 2017). In this study, RWC was reduced by approximately 12% overall during the water deficit periods (Fig. S8). This relatively small decrease in RWC is likely due to the fact that although water was reduced in the WS treatments, small amounts of water were delivered on a regular basis, as opposed to a long-term dry-down (de Silva *et al.*, 2012; Catoni *et al.*, 2017). In all species, RWC increased to control levels shortly after irrigation of the WS plants, as has been observed previously when an imposed water deficit did not severely reduce leaf RWC (Tognetti *et al.*, 1995). In this way, our water deficit treatment mimicked the type of stress encountered by trees in periods with reduced precipitation compared to wet periods (Kubiske & Abrams, 1991; Backes & Leuschner, 2000). This established that our experiments tested acclimation responses as they may arise in natural drought conditions, rather than leaf responses to rapid dehydration.

Some tree species are able to adjust osmotically in response to water deficit episodes (Ranney *et al.*, 1991; Wang & Stutte, 1992), whereas other species show no evidence of osmotic adjustment (OA) under water deficit conditions (Tschaplinski *et al.*, 1995). The observed maintenance of high RWC in WS leaves generally occurred without OA. In the 60% MWC treatment, there was no evidence for OA in any species (Fig. S9d–f). Redbud accumulates soluble carbohydrates in response to WS, but apparently do not osmotically adjust, as shown in the current and past (Griffin *et al.*, 2004) experiments. The only evidence we found of OA was in 40% MWC maple (Fig. S9e). This OA under only severe WS has been observed in *Fraxinus excelsior* (Guicherd *et al.*, 1997) and hybrid *Populus* genotypes (Gebre *et al.*, 1998). However, this did not affect the water status or growth of leaves, as both species produced smaller leaves (Figs S5, S7), and maple leaves still had reduced RWC despite the OA (Fig. S8b). Altogether, the growth and water relations response to WS was similar across silver maple, river birch, and eastern redbud leaves.

### Stomatal development plasticity in trees via different mechanisms to a common outcome

Stomatal closure in response to WS is a common response (Loewenstein & Pallardy, 1998; Bréda *et al.*, 2006). However, the plasticity of stomatal anatomy in leaves that emerge under water-deficit conditions is far less studied, especially in a manner that integrates different stomatal traits to show the final overall change in stomatal anatomy across the leaf epidermis. It is likely that this anatomical plasticity plays some role in acclimation to water deficit, since the molecular basis for stomatal development plasticity in response to water deficit has been established in some tree species (Hamanishi *et al.*, 2012; Viger *et al.*, 2016).

Reduced SPI in response to water deficit has been demonstrated in some tree species (Gindel, 1969; Camposeo *et al.*, 2011), but is not the typical response in trees (unpublished metaanalysis; Aasamaa *et al.*, 2001; Luo *et al.*, 2007; Eksteen *et al.* 2013) due to no anatomical changes occurring or the fact that the often-observed reduction in SS is not sufficient to have an impact on SPI (Aasamaa *et al.*, 2001; Luo *et al.*, 2007; Machado *et al.*, 2010; de Silva *et al.*, 2012; Hovenden *et al.*, 2012; Eksteen *et al.*, 2013; Catoni *et al.*, 2017). However, it is clear from our study that some tree species do respond to water deficit stress via stomatal development plasticity. In birch and redbud, we observed a common outcome of reduced SPI, but the basis of SPI plasticity differed between the species and stress severity. Under the 60% MWC treatment, birch stomata were smaller, whereas redbud leaves had fewer stomata. Under the 40% MWC treatment, redbud leaves instead had smaller stomata. Water-deficit-induced reductions in SD (Pääkkönen *et al.*, 1998; Silva, *et al.*, 2009; Camposeo *et al.*, 2011; Rajabpoor *et al.*, 2014) and SS (Luo *et al.*, 2007; Maes *et al.*, 2009) have been noted in other tree species. Many of these studies were conducted in dry regions and/or dry-adapted species, but in this study we show that these mechanisms of stomatal development plasticity also exist in temperate North American species adapted to more mesic environments.

The response to WS by maple leaves is more reflective of the broader literature on stomatal anatomy plasticity. This lack of stomatal development plasticity may be because maintenance of existing stomatal (and thus gas-exchange) capacity is advantageous for post-drought recovery, and because transient responses to water deficit, such as stomatal closure or solute accumulation, can be easily reversed, whereas anatomical changes to leaves are permanent. Still, the fact that there is a molecular basis and empirical data for stomatal anatomy plasticity in tree species suggests a potential acclimation/adaptive role.

The basis of SPI variation differs among species even under WW conditions. In WW birch and redbud leaves, higher SPI was a product of higher SD and SS (Fig. S21 a, c), but in maple leaves, higher SPI was due to larger stomata (Fig. S21b). Because SPI was correlated with stomatal frequency and size in WW and WS leaves, it appears that different tree species allocate a similar epidermal allocation of stomatal pore area (similar range of SPI in all three species, Fig. S21) differently via alteration of SS and/or SD. Silver maple leaves appear to have evolved to maintain a minimal range of small stomata while maximizing SPI via SD (Franks *et al.*, 2009). Maintaining small stomata as SPI variation is dependent on variation in SD would also minimize the cost associated with opening stomata (Spence *et al.*, 1986; Raven *et al.*, 2014), while maximizing the benefit (potential conductance or *g_smax_*) obtained by increasing SD (de Boer *et al*., 2016). Redbud WS leaves also had a relatively tight range of small stomata, but exhibited SPI variation via a much broader range of SD (Fig S21c).

It appears that the components of SPI are subject to independent mechanisms of plasticity, resulting in different anatomical mechanisms to a common outcome (Fig. 5). Firstly, passive control of guard cell size is evident in birch WS leaves because both cell types were smaller in WS leaves (birch, Figs 1, S13a, S15a, b) and pavement cell size and SS were positively correlated (Fig. S22a, c). Passive control of stomatal size, whereby guard cell dimensions are a function of cell turgor, probably result in a mechanical advantage frequently ascribed to smaller stomata (Spence *et al.*, 1986), and thus is favored under WS conditions. Under WW conditions, SS could be actively controlled to respond to other environmental factors, such as light.

**Fig. 5.**
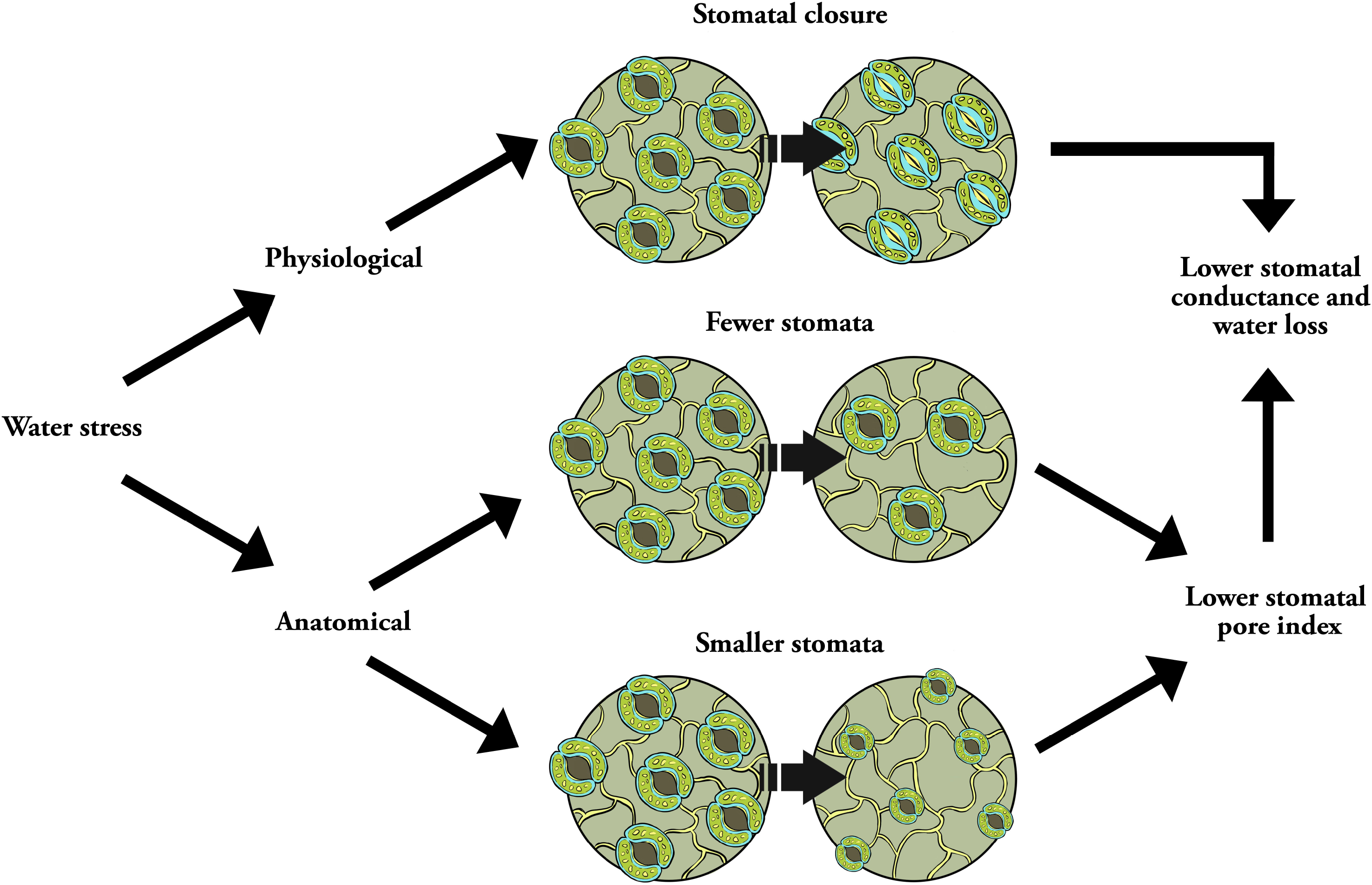
Schematic of the changes in stomatal anatomy observed in different tree species subjected to 60 or 40% MWC water deficit. The common outcome of changes in density (redbud, 60% MWC) or size (birch, 60% MWC; redbud, 40% MWC) was a reduction in the total stomatal area of water-stressed leaves, such that stomatal conductance was reduced. This outcome was also achieved in WS maple, albeit by stomatal closure instead of stomatal development plasticity.

Redbud WW and WS leaves exhibited differential coordination of stomatal and pavement cell size (Fig. S22c), raising the possibility of different mechanisms controlling stomatal size in response to changing environmental conditions. Previous work in *Arabidopsis* has demonstrated that pavement and guard cell size develop differently during leaf growth due to differential regulation of cell growth (Asl *et al.*, 2011).

The WS-induced changes in stomatal traits appear to be elicited by different mechanisms in the different species. Specifically, we observed that decreases in birch SI, SS, and SPI were correlated with decreased leaf RWC (Fig. S23). Additionally, since guard and pavement cells were smaller in birch leaves (Figs 1c, S13a, S15a, b), stomatal trait plasticity may be primarily turgor-driven in this species. This response in birch leaves may be due to a reduced RWC-induced accumulation of abscisic acid in drought-stressed leaves (Sack *et al.*, 2018). The sensitivity of stomatal and leaf development pathways to ABA accumulation may be more pronounced in certain species such as birch via variation in expression, copy number, or protein homology of the genes involved in ABA signaling and leaf cell development.

Vein density is often higher in more drought-tolerant species or genotypes, often coupled with reduced LA (Scoffoni *et al.*, 2011; Nardini *et al.*, 2012), which may enable leaf hydration during drought conditions, as embolized veins can be bypassed through additional venation (Sack *et al.*, 2008). Although our VD data was similar to prior values for related species (Sellin *et al.*, 2012; Uhl & Mosbrugger, 1999), VD was similar in WW and WS leaves (Figs 3, S18). In a variety of species, Aasamaa *et al*. (2001) also found VD to be similarly non-plastic. Maple leaves grown under 40% MWC were the only case in which VD increased in response to water deficit (Figs 3. S18b), possibly as part of the overall response in which there was no change in stomatal development or SPI. This may suggest that only a severe water deficit necessitates a change in water supply to leaves, but this response may itself be absent if leaf water demand is reduced by other anatomical plasticity such as lower SPI in redbud leaves. Fiorin *et al*. (2016) propose that the mean stomata-vein distance imposes a limit on the density of stomata that can be adequately supplied water by leaf venation. With this in mind, the reduction of leaf thickness (in the mild stress) and stomatal or vein plasticity may have been sufficient to keep leaf tissue and the associated stomata hydrated.

The coordination of stomatal and vein development has been described as the balance between water demand and supply in leaves (Brodribb & Jordan, 2011; Schneider *et al.*, 2017). To achieve a high rate of gas exchange, a high SD must be matched with high VD (Fiorin *et al.*, 2016), which has often been demonstrated in the correlation between SD and VD (Carins Murphy *et al.*, 2012; Carins Murphy *et al.*, 2016). In *T. ciliata,* this relationship was found to persist even after imposition of a low humidity treatment (Carins Murphy *et al.*, 2014). Although maple leaves did not exhibit stomatal development plasticity in response to WS, SD was nevertheless coordinated with VD in the range of SD across WW and WS leaves (Fig. S24b). Vein density also increased in 40% MWC maple leaves (Fig. S18b), and it is unlikely that this was simply due to smaller leaves under this treatment, as SD did not similarly increase (Figs 1a, S11b). Thus, WS maple leaves may produce additional venation to maintain the positive SD-VD correlation (Carins Murphy *et al.*, 2014). Furthermore, the SD-VD relationship was distinct in redbud and birch WS versus WW leaves (Fig. S24). Although SD-VD coordination has been proposed as a critical factor in the evolution of angiosperms (Boyce *et al.*, 2009; Zhang *et al.*, 2012), it is nonetheless absent in several woody angiosperm species (Torre *et al.*, 2003; Zhao *et al.*, 2016).

### Impact of stomatal development plasticity on leaf physiology

Water deficit results in stomatal closure, and thus reduced *g_s_* and *A* in leaves of river birch (Ranney *et al.*, 1991), eastern redbud (Abrams, 1988; Griffin *et al.*, 2004) and related maple species (Bauerle *et al.*, 2003). Stomatal closure is sometimes observed in conjunction with stomatal plasticity (Cavender-Bares *et al.*, 2007; Eksteen *et al.*, 2013). In a diverse panel of gymnosperms, ferns, and angiosperms, McElwain *et al.* (2016) showed that as the anatomical capacity for gas exchange increases, the operational rate of gas exchange also increases. However, in this study, gas exchange was not strongly dependent on stomatal anatomy. In birch and redbud, but not maple leaves, stomatal development was positively correlated with *A* and *g_s_* (Fig. S25). However, many other stomatal anatomical traits were not correlated with gas exchange, and especially critically, neither SPI nor *g_smax_* were correlated with *g_s_* (data not shown). The lack of a clear link between stomatal anatomy and gas exchange during drought conditions has previously been reported in a variety of species (Pääkkönen *et al.*, 1998; Carins Murphy *et al.*, 2014; Vieira *et al.*, 2017; Toscano *et al.*, 2018) Based on these reports and the present data set, we propose that among tree species, stomatal traits are only partially responsible for leaf physiology during water deficit episodes.

Plasticity in SPI was frequently not accompanied by plasticity in *g_smax_* (Figs 1a, 2), with only redbud leaves under 60% MWC exhibiting a decrease in both traits. Other instances of reduced SPI, such as birch leaves under 60% MWC and redbud leaves under 40% MWC, were not accompanied by a decrease in *g_smax_.* Reducing leaf SPI but maintaining *g_smax_* thus minimizes stomatal production and operating costs while maximizing carbon assimilation gains, which is especially important in water-stressed leaves. Moreover, maintenance of *g_smax_* would enable rapid return to normal *g_s_* after restoration of water-sufficient conditions, especially if a similar overall anatomy of water-stressed leaves now comprises smaller stomata that can be opened more easily. Because most species, including those from the present dataset, show operational *g_s_* as a very low fraction of *g_smax_*, (Fig. S26; McElwain *et al.*, 2016), the reduction of *g_smax_* in 60% MWC redbud leaves would have little to no impact on redbud recovery, and indeed this was observed in redbud following root-zone re-saturation (Fig. 4c, f).

Instead of a balance between stomatal costs and benefits, it may be the case that stomatal anatomy under water-deficit conditions is directed towards facilitating stomatal closure. None of the anatomical traits that control the dimensions and overall area of stomata for gas exchange (SD, SS, SPI, *g_smax_)* were correlated with operational *A* or *g_s_.* Instead, in all three species, the area coverage of stomata was correlated with the degree of stomatal closure observed during the stress period (Fig. S27). Thus, lower SPI reduced the degree of stomatal closure necessary during the WS period. Although not typically discussed in these terms, we propose that stomatal developmental inhibition is aimed towards achieving minimum *g_s_*, without constraining maximum *g_s_*.

## Conclusions

A common outcome of reduced SPI is achieved by some North American tree species via different mechanisms in response to water deficit (Fig. 5). These species and treatment-level differences illustrate the importance of reporting all stomatal traits in leaf anatomical plasticity studies. For instance, although SS or SD was reduced in some *Eucalyptus grandis* clones under certain water-deficit treatments, SPI calculated from these values was almost always unchanged (Eksteen *et al.*, 2013). A similar effect was observed in certain *Prunus dulcis* ecotypes: reduced SPI was the result of either smaller or fewer stomata, so presenting these traits in isolation would have missed this phenotypic plasticity (Camposeo *et al.*, 2011). In all three species across both treatment levels, examining stomatal frequency and size, as well as the combinations of these traits via SPI and *g_smax_*, was critical to the conclusions drawn. Had SD or SS been examined in isolation, the different mechanisms of plasticity between birch and redbud at 60% MWC would have not been revealed. Similarly, the differential response in redbud leaves grown at 60% or 40% MWC would have also been missed. We could also deduce that reduced *g_s_* in WS maple (and the fourth leaf of WS birch) was due primarily to reduced stomatal aperture, since there were no changes in the total coverage of stomata in these leaves, despite the smaller stomata produced in these leaves under the 40% MWC treatment (Figs 1c, S13).

## Acknowledgements

We thank James McKenna for plant material; Nathan Deppe and Dan Little for greenhouse assistance; Mike Gosney, Amanda Ávila Cardoso, and Huangai Bi for assistance with data collection; Robert Heath for assistance with leaf growth analysis; Scott McAdam and Chris Oakley for suggestions on methodology and manuscript advice; Peter Goldsbrough for helpful edits to the manuscript; Tom Kronewitter of Purdue University Agricultural Communication for the production of Figure 5; and the Purdue University Center for Plant Biology for funding.

## Author contributions

NAM and MVM designed the experiment; NAM and SFL performed all greenhouse work and data collection; NAM and MVM analyzed the data; NAM, SFL, and MVM wrote the manuscript.

## Supporting information

**Figure S1.** Environmental conditions throughout the experiment.

**Figure S2.** Media water content throughout the experiment.

**Figure S3.** Schematic of leaves used for data collection in the 60% MWC treatment.

**Table S1.** Description of measurements made on different leaves during both experiments.

**Figure S4.** Leaf growth dynamics in the 60% MWC treatment.

**Figure S5.** Plasticity of tree height and leaf area in both experiments.

**Figure S6.** Height of trees after both treatments of WS.

**Figure S7.** Leaf area of individual leaves after both WS treatments.

**Figure S8.** Relative water content of individual leaves in both WS treatments.

**Figure S9.** Leaf osmotic potential and osmotic potential at full turgor after both WS treatments.

**Figure S10.** Specific leaf weight of leaves after both WS experiments.

**Figure S11.** Stomatal density of individual leaves after both experiments.

**Figure S12.** Stomatal index of individual leaves after both experiments.

**Figure S13.** Stomatal size of individual leaves after both experiments.

**Figure S14.** Stomatal length and width of individual leaves after both experiments.

**Figure S15.** Collected plasticity and individual leaf pavement cell size after both experiments.

**Figure S16.** Collected plasticity and individual leaf pavement cell density after both experiments.

**Figure S17.** Stomatal pore index of individual leaves after both WS experiments.

**Figure S18.** Vein density in leaves 3 and 4 measured after development under well-watered or water-stressed (WS, 60 or 40% MWC) conditions.

**Figure S19.** Transpiration rate and whole-leaf transpiration in leaves during and after the 60% MWC water deficit.

**Figure S20.** Stomatal conductance, net CO_2_ assimilation, transpiration rate, and whole-leaf transpiration during the 40% MWC water deficit.

**Figure S21.** The relationship between stomatal pore index and stomatal size, density, and index across both experiments.

**Figure S22.** Relationship between pavement cell size and stomatal size across both experiments.

**Figure S23.** Relationship between relative water content and stomatal traits across both experiments.

**Figure S24.** Relationship between stomatal and vein density across both experiments.

**Figure S25.** Relationship between stomatal index and gas exchange across both experiments.

**Figure S26.** Changes in stomatal conductance as a % of *g_smax_* in the fourth leaf that developed under water-stressed conditions (60% MWC), along with well-watered controls over time.

**Figure S27.** Relationship between stomatal pore index and degree of stomatal closure in WS plants.

## References

Aasamaa K, Sõber A, Rahi M. 2001. Leaf anatomical characteristics associated with shoot hydraulic conductance, stomatal conductance and stomatal sensitivity to changes of leaf water status in temperate deciduous trees. Australian Journal of Plant Physiology 28: 765–774.

Abrams MD. 1988. Genetic variation in leaf morphology and plant and tissue water relations during drought in *Cercis canadensis* L. Forest Science 34: 200–207.

Andriankaja M, Dhondt S, De Bodt S, Vanhaeren H, Coppens F, De Milde L, Mühlenbrock P, Skirycz A, Gonzalez N, Beemster GTS, et al. 2012. Exit from proliferation during leaf development in *Arabidopsis thaliana:* a not-so-gradual process. Developmental Cell 22: 64–78.

Asl LK, Dhondt S, Boudolf V, Beemster GTS, Beeckman T, Inzé D, Govaerts W, De Veylder L. 2011. Model-based analysis of Arabidopsis leaf epidermal cells reveals distinct division and expansion patterns for pavement and guard cells. Plant Physiology 156: 2172 – 2183.

Aspelmeier S, Leuschner C. 2006. Genotypic variation in drought response of silver birch *(Betula pendula* Roth): leaf and root morphology and carbon partitioning. Trees 20: 42–52.

Backes K, Leuschner C. 2000. Leaf water relations of competitive *Fagus sylvatica* and *Quercus petraea* trees during 4 years differing in soil drought. Canadian Journal of Forest Research 30: 335–346.

Baerenfaller K, Massonnet C, Walsh S, Baginsky S, Bühlmann P, Hennig L, Hirsch– Hoffmann M, Howell KA, Kahlau S, Radziejwoski et al. 2012. Systems-based analysis of Arabidopsis leaf growth reveals adaptation to water deficit. Molecular Systems Biology 8: 606.

Bauerle WL, Dudley JB, Grimes LW. 2003. Genotypic variability in photosynthesis, water use, and light absorption among red and Freeman maple cultivars in response to drought stress. Journal of the American Society for Horticultural Science 128: 337–342.

Boyce CK, Brodribb TJ, Feild TS, Zwieniecki MA. 2009. Angiosperm leaf vein evolution was physiologically and environmentally transformative. Proceedings of the Royal Society B 276: 1771–1776.

Bréda N, Huc R, Granier A, Dreyer E. 2006. Temperate forest trees and stands under severe drought: a review of ecophysiological responses, adaptation processes and long-term consequences. Annals of Forest Science 63: 625–644.

Breshears DD, Myers OB, Meyer CW, Barnes FJ, Zou CB, Allen CD, McDowell NG, Pockman WT. 2009. Tree die-off in response to global change-type drought: mortality insights from a decade of plant water potential measurements. Frontiers in Ecology and the Environment 7: 185–189.

Brodribb TJ, Jordan GJ. 2011. Water supply and demand remain balanced during leaf acclimation of *Nothofagus cunninghamii* trees. New Phytologist 192: 437–448.

Camposeo S, Palasciano M, Vivaldi GA, Godini A. 2011. Effect of increasing climatic water deficit on some leaf and stomatal parameters of wild and cultivated almonds under Mediterranean conditions. Scientia Horticulturae 127: 234–241.

Carins Murphy MR, Jordan GJ, Brodribb TJ. 2012. Differential leaf expansion can enable hydraulic acclimation to sun and shade. Plant, Cell and Environment 35: 1407–1418.

Carins Murphy MR, Jordan GJ, Brodribb TJ. 2014. Acclimation to humidity modifies the link between leaf size and the density of veins and stomata. Plant, Cell and Environment 37: 124–131.

Carins Murphy MR, Jordan GJ, Brodribb TJ. 2016. Cell expansion not cell differentiation predominantly co-ordinates veins and stomata within and among herbs and woody angiosperms grown under sun and shade. Annals of Botany 118: 1127–1138.

Casson SA, Franklin KA, Gray JE, Grierson CS, Whitelam GC, Hetherington AM. 2009. Phytochrome B and *PIF4* regulate stomatal development in response to light quantity. Current Biology 19: 229–234.

Catoni R, Gratani L, Bracco F, Granata MU. 2017. How water supply during leaf development drives water stress response in *Corylus avellana* saplings. Scientia Horticulturae 214: 122–132.

Cavender-Bares J, Sack L, Savage J. 2007. Atmospheric and soil drought reduce nocturnal conductance in live oaks. Tree Physiology 27: 611–620.

de Boer HJ, Price CA, Wagner-Cremer, Dekker SC, Franks PJ, Veneklaas EJ. 2016. Optimal allocation of leaf epidermal area for gas exchange. New Phytologist 210: 1219–1228.

de Silva NDG, Cholewa E, Ryser P. 2012. Effects of combined drought and heavy metal stresses on xylem structure and hydraulic conductivity in red maple *(Acer rubrum* L.). Journal of Experimental Botany 63: 5957–5966.

Donselman HM, Flint HL. 1982. Genecology of eastern redbud *(Cercis canadensis)*. Ecology 63: 962–971.

Dow GJ, Bergmann CD, Berry JA. 2014. An integrated model of stomatal development and leaf physiology. New Phytologist 201: 1218–1226.

Drake PL, Froend RH, Franks PJ. 2013. Smaller, faster stomata: scaling of stomatal size, rate of response, and stomatal conductance. Journal of Experimental Botany 64: 495–505.

Dunlap JM, Stettler RF. 2001. Variation in leaf epidermal and stomatal traits of *Populus trichocarpa* from two transects across the Washington Cascades. Canadian Journal of Botany 79: 528–536.

Eksteen AB, Grzeskowiak V, Jones NB, Pammenter NW. 2013. Stomatal characteristics of *Eucalyptus grandis* clonal hybrids in response to water stress. Southern Forests 75: 105–111.

Fiorin L, Brodribb TJ, Anfodillo T. 2016. Transport efficiency through uniformity: organization of veins and stomata in angiosperm leaves. New Phytologist 209: 216–227.

Franks PJ, Farquhar GD. 2001. The effect of exogenous abscisic acid on stomatal development, stomatal mechanics, and leaf gas exchange in *Tradescantia virginiana*. Plant Physiology 125: 935–942.

Franks PJ, Beerling DJ. 2009. Maximum leaf conductance driven by CO2 effects on stomatal size and density over geologic time. Proceedings of the National Academy of Sciences, USA 106: 10343–10347.

Franks PJ, Drake PL, Beerling DJ. 2009. Plasticity in maximum stomatal conductance constrained by negative correlation between stomatal size and density: an analysis using *Eucalyptus globulus*. Plant, Cell & Environment 32: 1737–1748.

Fritsch PW, Nowell CF, Leatherman LST, Gong W, Cruz BC, Burge DO, Delgado-Salinas A. 2018. Leaf adaptations and species boundaries in North American *Cercis:* implications for the evolution of dry floras. American Journal of Botany 105: 1577–1594.

Gebre GM, Tschaplinski TJ, Tuskan GA, Todd DE. 1998. Clonal and seasonal differences in leaf osmotic potential and organic solutes of five hybrid poplar clones grown under field conditions. Tree Physiology 18: 645–652.

Geisler M, Nadeau J, Sack FD. 2000. Oriented asymmetric divisions that generate the stomatal spacing pattern in Arabidopsis are disrupted by the *too many mouths* mutation. The Plant Cell 12: 2075–2086.

Giday H, Kjaer KH, Fanourakis D, Ottosen C-O. 2013. Smaller stomata require less severe leaf drying to close: a case study in *Rosa hybrida*. Journal of Plant Physiology 170: 1309–1316.

Gindel I. 1969. Stomatal number and size as related to soil moisture in tree xerophytes in Israel. Ecology 50: 263–267.

Gitlin AR, Sthultz CM, Bowker MA, Stumpf S, Paxton KL, Kennedy K, Muñoz A, Bailey JK, Whitham TG. 2006. Mortality gradients within and among dominant plant populations as barometers of ecosystem change during extreme drought. Conservation Biology 20: 1477–1486.

Griffin JJ, Ranney TG, Pharr DM. 2004. Heat and drought influence photosynthesis, water relations, and soluble carbohydrates of two ecotypes of redbud (*Cercis canadensis*). Journal of the American Society for Horticultural Science 129: 497–502.

Guicherd P, Peltier JP, Gout E, Bligny R, Marigo G. 1997. Osmotic adjustment in *Fraxinus excelsior* L.: malate and mannitol accumulation in leaves under drought conditions. Trees 11: 157–161.

Guo X, Luo Y-J, Xu Z-W, Li M-Y, Guo W-H. 2019. Response strategies of *Acer davidii* to varying light regimes under different water conditions. Flora 257: 151423.

Hamanishi ET, Thomas BR, Campbell MM. 2012. Drought induces alterations in the stomatal development program in *Populus*. Journal of Experimental Botany 63: 4959–4971.

Harb A, Krishnan A, Ambavaram MMR, Pereira A. 2010. Molecular and physiological analysis of drought stress in Arabidopsis reveals early responses leading to acclimation in plant growth. Plant Physiology 154: 1254–1271.

Hovenden MJ, Vander Schoor JK. 2012. Soil water potential does not affect leaf morphology or cuticular characters important for palaeo-environmental reconstructions in southern beech, *Nothofagus cunninghamii* (Nothofagaceae). Australian Journal of Botany 60: 87–95.

Hovenden MJ, Vander Schoor JK, Osanai Y. 2012. Relative humidity has dramatic impacts on leaf morphology but little effect on stomatal index or density in *Nothofagus cunninghamii* (Nothofagaceae). Australian Journal of Botany 60: 700–706.

Jin Z, Ainsworth EA, Leakey ADB, Lobell DB. 2018. Increasing drought and diminishing benefits of elevated carbon dioxide for soybean yields across the US Midwest. Global Change Biology 24: e522–e533.

Kang C-Y, Lian H-L, Wang F-F, Huang J-R, Yang H-Q. 2009. Cryptochromes, phytochromes, and COP1 regulate light-controlled stomatal development in *Arabidopsis*. The Plant Cell 21: 2624–2641.

Kang J, Dengler N. 2004. Vein pattern development in adult leaves of *Arabidopsis thaliana*. International Journal of Plant Sciences 165: 231–242.

Kilian J, Whitehead D, Horak J, Wanke D, Weinl S, Batistic O, D’Angelo C, Bornberg-Bauer E, Kudla J, Harter K. 2007. The AtGenExpress global stress expression data set: protocols, evaluation and model data analysis of UV–B light, drought and cold stress responses. Plant Journal 50: 347–363.

Kitao M, Lei TT, Koike T, Tobita H, Maruyama Y. 2003. Higher electron transport rate observed at low intercellular CO2 concentration in long-term drought-acclimated leaves of Japanese mountain birch *(Betula ermanii)*. Physiologia Plantarum 118: 406–413.

Kleczewski NM, Herms DA, Pierluigi B. 2012. Nutrient and water availability alter belowground patterns of biomass allocation, carbon partitioning, and ectomycorrhizal abundance in *Betula nigra*. Trees 26: 525–533.

Klos RJ, Wang GG, Bauerle WL, Rieck JR. 2009. Drought impact on forest growth and mortality in the southeast USA: an analysis using Forest Health and Monitoring data. Ecological Applications 19: 699–708.

Kubiske ME, Abrams MD. 1991. Rehydration effects on pressure-volume relationships in four temperate woody species: variability with site, time of season and drought conditions. Oecologia 85: 537–542.

Kumari A, Jewaria PK, Bergmann DC, Kakimoto T. 2014. Arabidopsis reduces growth under osmotic stress by decreasing SPEECHLESS protein. Plant & Cell Physiology 55: 2037–2046.

Li L, Wang X, Niu J, Cui J, Zhang Q, Wan W, Liu B. 2017. Effects of elevated atmospheric O3 concentrations on early and late leaf growth and elemental contents of *Acer truncatum* Bung under mild drought. Acta Ecologica Sinica 37: 31–34.

Liu C, He N, Zhang J, Li Y, Wang Q, Sack L, Yu G. 2018. Variation of stomatal traits from cold temperate to tropical forests and association with water use efficiency. Functional Ecology 32: 20–28.

Loewenstein NJ, Pallardy SG. 1998. Drought tolerance, xylem sap abscisic acid and stomatal conductance during soil drying: a comparison of canopy trees of three temperate deciduous angiosperms. Tree Physiology 18: 431–439.

Ludlow AE. 1991. *Ochna pulchra* Hook: leaf growth and development related to photosynthetic activity. Annals of Botany 68: 527–540.

Luo HJ, Zheng ZB, Luo S, Pan YS, Liu XH. 2007. Changes in leaf characters of loquat under repeated drought stresses. Acta Horticulturae 750: 417–422.

Ma P, Bai T-H, Ma F-W. 2015. Effects of progressive drought on photosynthesis and partitioning of absorbed light in apple trees. Journal of Integrative Agriculture 14: 681–690.

Machado AV, de Souza TV, Paulilo MTS, Santos M. 2010. Response of a woody species from Atlantic rain forest, *Hedyosmum brasiliense* Mart. ex Miq. (Chloranthaceae), submitted to water stress. Insula 39: 01–13.

Maes WH, Achten WMJ, Reubens B, Raes D, Samson R, Muys B. 2009. Plant-water relationships and growth strategies of *Jatropha curcas* L. seedlings under different levels of drought stress. Journal of Arid Environments 73: 877–884.

McDowell NG. 2011. Mechanisms linking drought, hydraulics, carbon metabolism, and vegetation mortality. Plant Physiology 155: 1051–1059.

McElwain JC, Chaloner WG. 1995. Stomatal density and index of fossil plants track atmospheric carbon dioxide in the Palaeozoic. Annals of Botany 76: 389–395.

McElwain JC, Yiotis C, Lawson T. 2016. Using modern plant trait relationships between observed and theoretical maximum stomatal conductance and vein density to examine patterns of plant macroevolution. New Phytologist 209: 94–103.

McKown AD, Guy RD, Quamme L, Klápštč J, La Mantia J, Constabel CP, El-Kassaby YA, Hamelin RC, Zifkin M, Azam MS. 2014. Association genetics, geography and ecophysiology link stomatal patterning in *Populus trichocarpa* with carbon gain and disease resistance trade-offs. Molecular Ecology 23: 5771–5790.

Nardini A, Pedà G, La Rocca N. 2012. Trade-offs between leaf hydraulic capacity and drought vulnerability: morpho-anatomical bases, carbon costs and ecological consequences. New Phytologist 196: 788–798.

Pääkkönen E, Vahala J, Pohjola M, Holopainen T, Kärenlampi L. 1998. Physiological, stomatal and ultrastructural ozone responses in birch *(Betula pendula* Roth.) are modified by water stress. Plant, Cell and Environment 21: 671–684.

Pearce DW, Millard S, Bray DF, Rood SB. 2006. Stomatal characteristics of riparian poplar species in a semi-arid environment. Tree Physiology 26: 211–218.

Ponce-Campos GE, Moran MS, Huete A, Zhang Y, Bresloff C, Huxman TE, Eamus D, Bosch DD, Buda AR, Gunter SA, et al. 2013. Ecosystem resilience despite large-scale altered hydroclimatic conditions. Nature 494: 349–352.

Possen BJHM, Oksanen E, Rousi M, Ruhanen H, Ahonen V, Tervahauta A, Heinonen J, Heiskanen J, Kärenlampi S, Vapaavuori E. 2011. Adaptability of birch *(Betula pendula* Roth) and aspen *(Populus tremula* L.) genotypes to different soil moisture conditions. Forest Ecology and Management 262: 1387–1399.

Rajabpoor S, Kiani S, Sorkheh K, Tavakoli F. 2014. Changes induced by osmotic stress in the morphology, biochemistry, physiology, anatomy and stomatal parameters of almond species *(Prunus* L. spp.) grown *in vitro*. Journal of Forestry Research 25: 523–534.

Ranney TG, Bassuk NL, Whitlow TH. 1991. Osmotic adjustment and solute constituents in leaves and roots of water-stressed cherry (*Prunus*) trees. Journal of the American Society for Horticultural Science 116: 684–688.

Raven JA. 2014. Speedy small stomata? Journal of Experimental Botany 6: 1415–1424.

Rawson HM, Craven CL. 1975. Stomatal development during leaf expansion in tobacco and sunflower. Australian Journal of Botany 23: 253–261.

Reddy AR, Chaitanya KV, Vivekanandan M. 2004. Drought-induced responses of photosynthesis and antioxidant metabolism in higher plants. Journal of Plant Physiology 161: 1189–1202.

Reyer CPO, Leuzinger S, Rammig A, Wolf A, Bartholomeus RP, Bonfante A, de Lorenzi F, Dury M, Gloning P, Abou Jaoudé R, et al. 2013. A plant’s perspective of extremes: terrestrial plant responses to changing climatic variability. Global Change Biology 19: 75–89.

Ruehr NK, Offermann CA, Gessler A, Winkler JB, Ferrio JP, Buchmann N, Barnard RL. 2009. Drought effects on allocation of recent carbon: from beech leaves to soil CO2 efflux. New Phytologist 184: 950–961.

Sack L, Cowan PD, Jaikumar N, Holbrook NM. 2003. The ‘hydrology’ of leaves: coordination of structure and function in temperate woody species. Plant, Cell and Environment 26: 1343–1356.

Sack L, Dietrich EM, Streeter CM, Sánchez-Gómez D, Holbrook NM. 2008. Leaf palmate venation and vascular redundancy confer tolerance of hydraulic disruption. Proceedings of the National Academy of Sciences, USA 105: 1567–1572.

Sack L, John GP, Buckley TN. 2018. ABA accumulation in dehydrating leaves is associated with decline in cell volume, not turgor pressure. Plant Physiology 176: 489–493.

Schneider JV, Habersetzer J, Rabenstein R, Wesenberg J, Wesche K, Zizka G. 2017. Water supply and demand remain coordinated during breakdown of the global scaling relationship between leaf size and major vein density. New Phytologist 214: 473–486.

Scoffoni C, Rawls M, McKown A, Cochard H, Sack L. 2011. Decline of leaf hydraulic conductance with dehydration: relationship to leaf size and venation architecture. Plant Physiology 156: 832–843.

Sellin A, Õunapuu E, Kaurilind E, Alber M. 2012. Size-dependent variability of leaf and shoot hydraulic conductance in silver birch. Trees 26: 821–831.

Skirycz A, Claeys H, De Bodt S, Oikawa A, Shinoda S, Andriankaja M, Maleux K, Eloy NB, Coppens F, Yoo S-D, et al. 2011. Pause-and-stop: the effects of osmotic stress on cell proliferation during early leaf development in *Arabidopsis* and a role for ethylene signaling in cell cycle arrest. The Plant Cell 23: 1876–1888.

Silva EC, Nogueira RJMC, Vale FHA, de Araújo FP, Pimenta MA. 2009. Stomatal changes induced by intermittent drought in four umbu tree genotypes. Brazilian Journal of Plant Physiology 21: 33–42.

Sobrado MA. 2007. Relationship of water transport to anatomical features in the mangrove *Laguncularia racemosa* grown under contrasting salinities. New Phytologist 173: 584–591.

Spence RD, Wu H, Sharpe PJH, Clark KG. 1986. Water stress effects on guard cell anatomy and the mechanical advantage of the epidermal cells. Plant, Cell and Environment 9: 197–202.

Syvertsen JP, Lloyd J, McConchie C, Kriedemann PE, Farquhar GD. 1995. On the relationship between leaf anatomy and CO_2_ diffusion through the mesophyll of hypostomatous leaves. Plant, Cell and Environment 18: 149–157.

Tabata A, Ono K, Sumida A, Hara T. 2010. Effects of soil water conditions on the morphology, phenology, and photosynthesis of *Betula ermanii* in the boreal forest. Ecology Research 25: 823–835.

Tipton JL, White M. 1995. Differences in leaf cuticle structure and efficacy among Eastern redbud and Mexican redbud phenotypes. Journal of the American Society for Horticultural Science 120: 59–64.

Tognetti R, Johnson JD, Michelozzi M. 1995. The response of European beech *(Fagus sylvatica* L.) seedlings from two Italian populations to drought and recovery. Trees 9: 348–354.

Torre S, Fjeld T, Gislerød HR, Moe R. 2003. Leaf anatomy and stomatal morphology of greenhouse roses grown at moderate or high air humidity. Journal of the American Society of Horticultural Science 128: 598–602.

Toscano S, Ferrante A, Tribulato A, Romano D. 2018. Leaf physiological and anatomical responses of Lantana and Ligustrum species under different water availability. Plant Physiology and Biochemistry 127: 380–392.

Tripathi P, Rabara RC, Reese RN, Miller MA, Rohila JS, Subramanian S, Shen QJ, Morandi D, Bücking H, Shulaev V, et al. 2016. A toolbox of genes, proteins, metabolites and promoters for improving drought tolerance in soybean includes the metabolite coumestrol and stomatal development genes. BMC Genomics 17: 102.

Tschaplinski TJ, Stewart DB, Norby RJ. 1995. Interactions between drought and elevated CO_2_ on osmotic adjustment and solute concentrations of tree seedlings. New Phytologist 131: 169–177.

Uhl D, Mosbrugger V. 1999. Leaf venation density as a climate and environmental proxy: a critical review and new data. Palaeogeography, Palaeoclimatology, Palaeoecology 149: 15–26.

Vieira EA, Silva MDG, Moro CF, Laura VA. 2017. Physiological and biochemical changes attenuate the effects of drought on the Cerrado species *Vatairea macrocarpa* (Benth.) Ducke. Plant Physiology and Biochemistry 115: 472–483.

Viger M, Smith HK, Cohen D, Dewoody J, Trewin H, Steenackers M, Bastien C, Taylor G. 2016. Adaptive mechanisms and genomic plasticity for drought tolerance identified in European black poplar *(Populus nigra* L.). Tree Physiology 36: 909–928.

Wang Z, Stutte GW. 1992. The role of carbohydrates in active osmotic adjustment in apple under water stress. Journal of the American Society for Horticultural Science 117: 816–823.

Woodall GS, Dodd IC, Stewart GR. 1998. Contrasting leaf development within the genus *Syzygium*. Journal of Experimental Botany 49: 79–87.

Woodward FI. 1987. Stomatal numbers are sensitive to increases in CO_2_ from pre-industrial levels. Nature 327: 617–618.

Wu Z, Dijkstra P, Koch GW, Peñuelas J, Hungate BA. 2011. Responses of terrestrial ecosystems to temperature and precipitation change: a meta-analysis of experimental manipulation. Global Change Biology 17: 927–942.

Yoo CY, Mano N, Finkler A, Weng H, Day IS, Reddy ASN, Poovaiah BW, Fromm H, Hasegawa PM, Mickelbart MV. 2019. A Ca^2+^/CaM-regulated transcriptional switch modulates stomatal development in response to water deficit. Scientific Reports 9: 12282.

Yoo CY, Pence HE, Jin JB, Miura K, Gosney MJ, Hasegawa PM, Mickelbart MV. 2010. The *Arabidopsis* GTL1 transcription factor regulates water use efficiency and drought tolerance by modulating stomatal density via transrepression of *SDD1*. The Plant Cell 22: 4128–4141.

Zhang S-B, Guan Z-J, Sun M, Zhang J-J, Cao K-F, Hu H. 2012. Evolutionary association of stomatal traits with leaf vein density in *Paphiopedilum*, Orchidaceae. PLoS One 7: e40080.

Zhao W-L, Chen Y-J, Brodribb TJ, Cao K-F. 2016. Weak co-ordination between vein and stomatal densities in 105 angiosperm tree species along altitudinal gradients in Southwest China. Functional Plant Biology 43: 1126–1133.

