## Supplemental data for "Divergent strategies to reduce stomatal pore index during water deficit in perennial angiosperms"

### *New Phytologist* Supporting Information

Article acceptance date:

The following Supporting Information is available for this article:

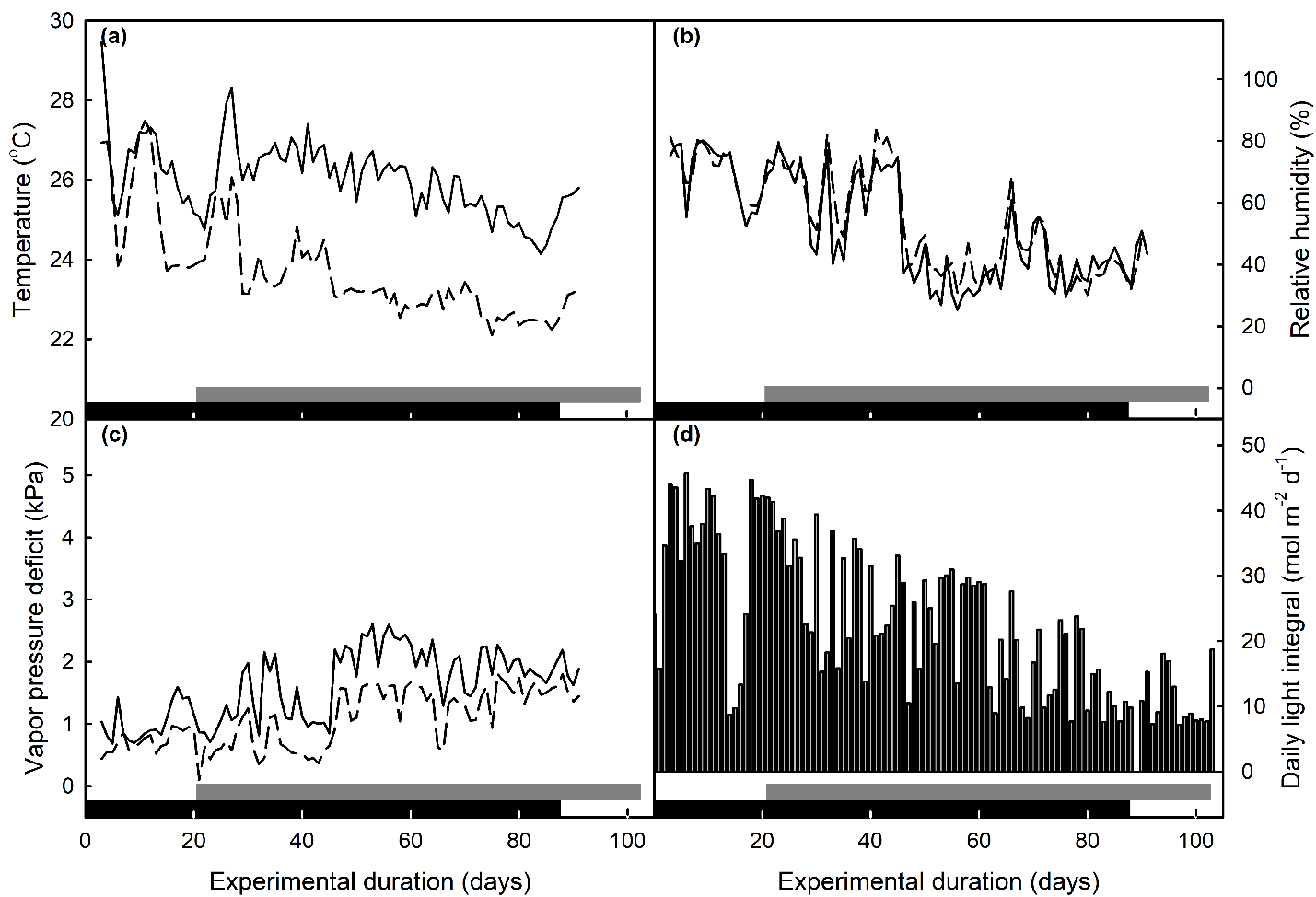

**Figure S1.** Average day (solid lines) and night (dashed lines) air temperature (a), relative humidity (b), vapor pressure deficit (c), and daily light integral (d) during experiments 1 (60% MWC, black bar on x-axis) and 2 (40% MWC, grey bar on x-axis). Data in panels a, b, and c are the average of two HOBO devices among the trees in the greenhouse, and daily light integral (d) was quantified with a weather station outside the greenhouse.

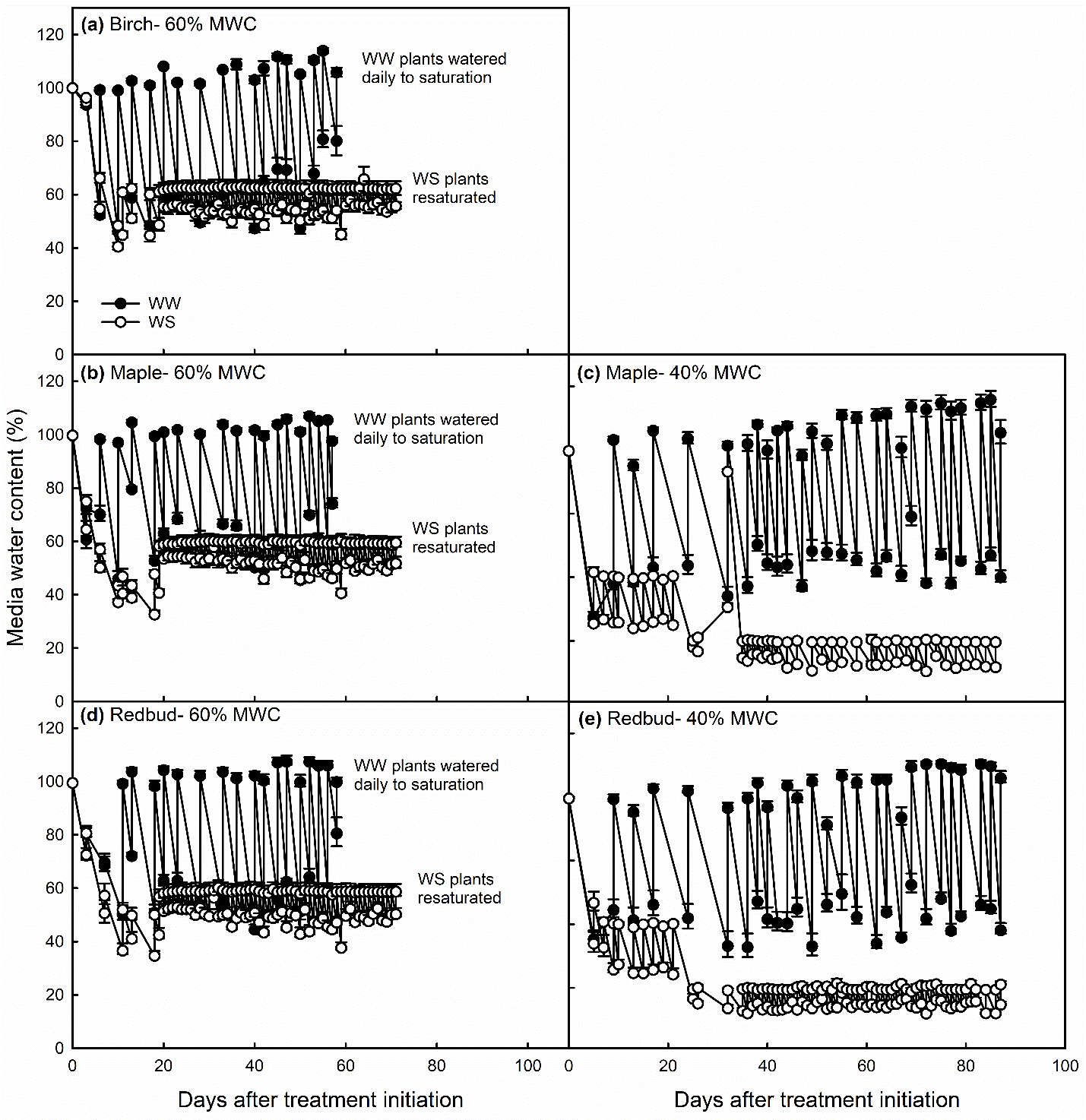

**Figure S2**. Media water content (MWC) in well-watered (WW, ●) and water-stressed (WS, ○) birch (a), maple (b), and redbud (d) during the 60% MWC treatment (*n =* 4–6), and maple (c) and redbud (e) during the 40% MWC treatment (*n =* 6–8). Data are means ± SE.

1

2

4

3

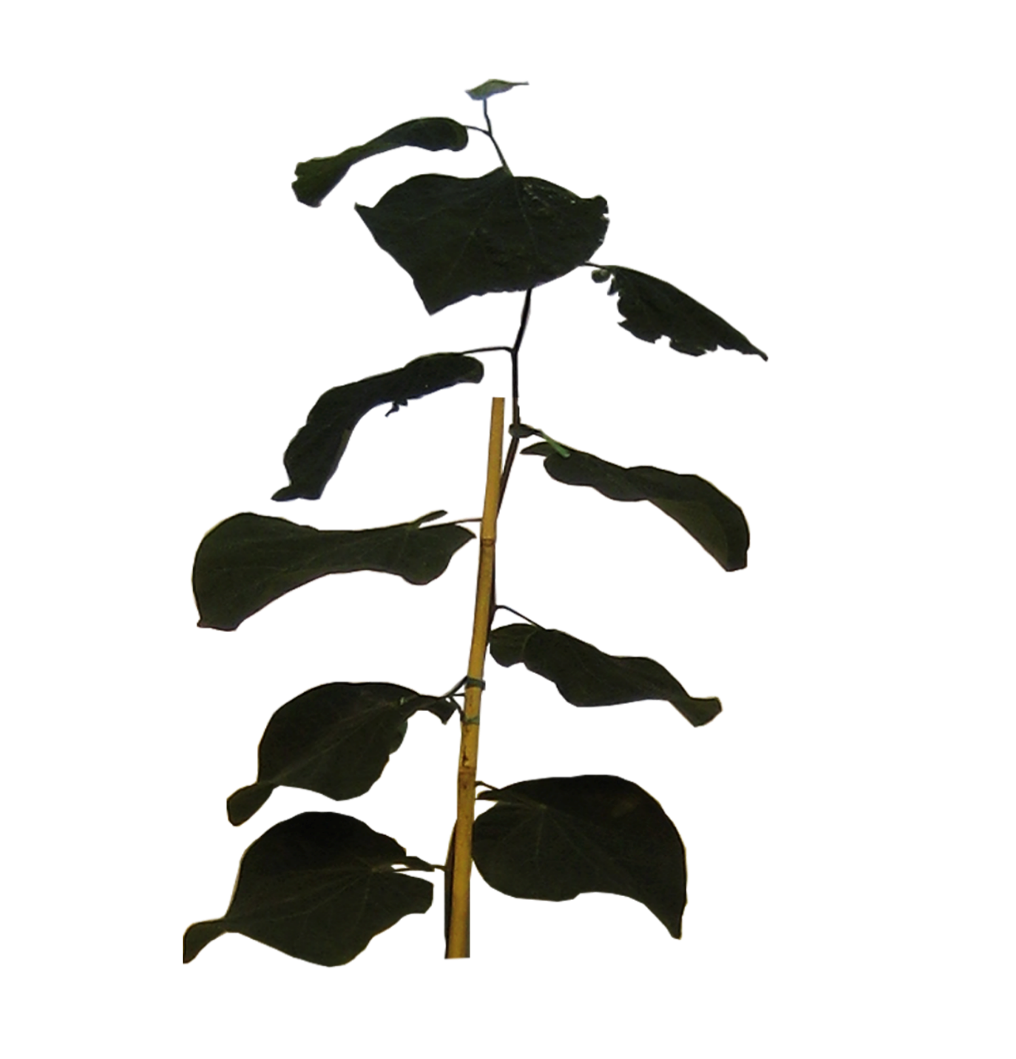

**Figure S3.** Leaves used for data collection, shown here for redbud (the same numbering scheme was used for maple). Leaf 1 was the first leaf to emerge after the target media water content (60% or 40%) was reached. In the first experiment, leaves 2, 3, and 4 were used and in the second experiment, leaf 1 was used for all measurements (Table S1). Because birch trees had multiple branches, a number of leaves equivalent to leaves 2 to 4 at the top of the canopy were used.

| **Table S1.** Description of measurements made on different leaves during both experiments. | | | |
| --- | --- | --- | --- |
| Treatment  (% MWC) | Leaf | Days after treatment initiation | Measurements |
| 60 | 2 | 60 | Epidermal traits, leaf area (LA), osmotic potential (Ψ_π_), osmotic potential at full turgor (Ψ_π100_), relative water content (RWC) |
|  | 3 | 61 | Epidermal traits, LA, vein density^†^, Ψ_π_, Ψ_π100_, RWC, gas exchange |
|  | 4 | 75−88 | Gas exchange |
|  |  | 82 | Redbud: Epidermal traits, LA, vein density, Ψ_π_, Ψ_π100_, RWC, gas exchange |
|  |  | 88 | Maple and birch: Epidermal traits, LA, vein density, Ψ_π_, Ψ_π100_, RWC, gas exchange |
| 40 | 1 | 82 | Epidermal traits, LA, vein density, Ψ_π_, Ψ_π100_, RWC, gas exchange |
| ^†^ Vein density was not measured on birch leaves on day 61. | | | |

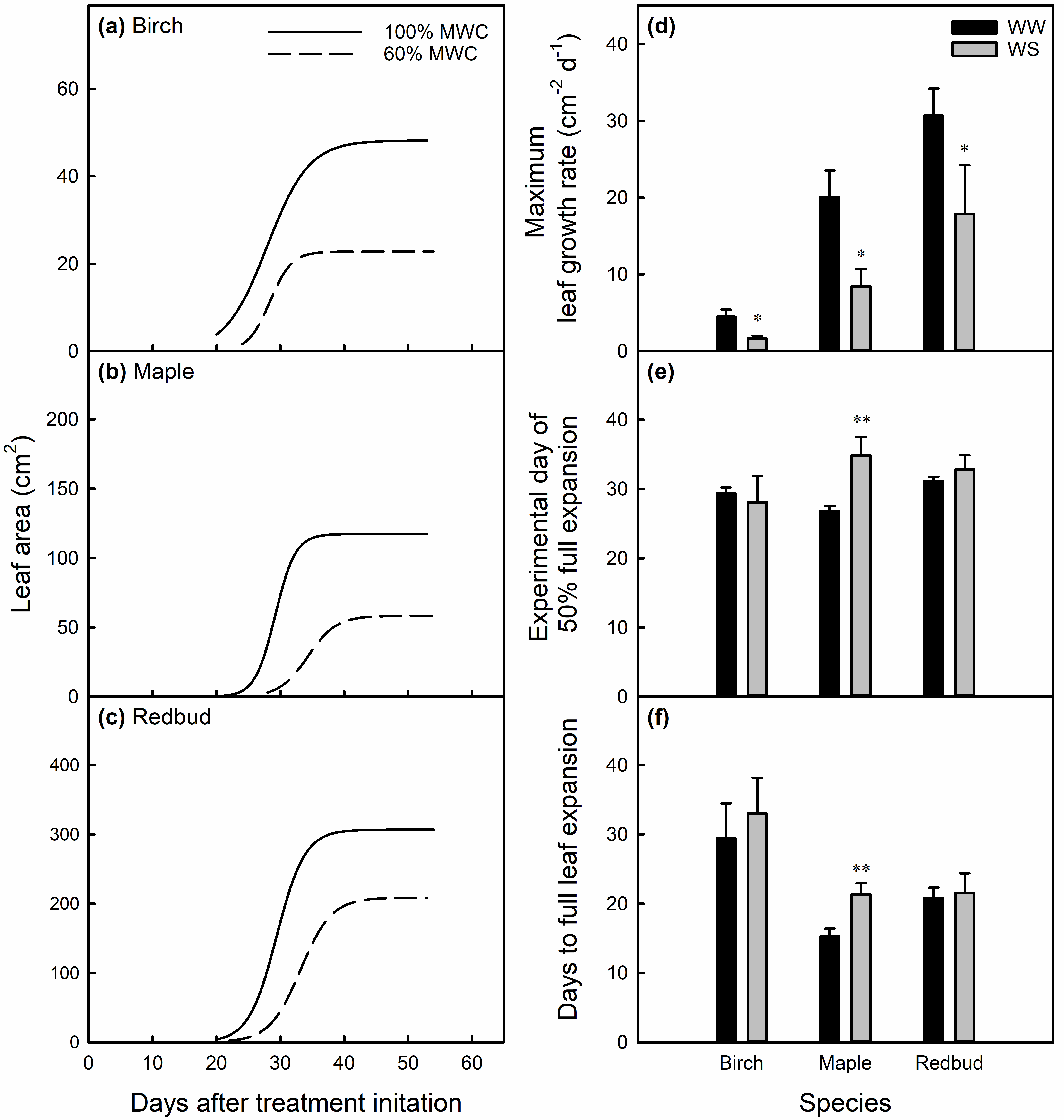

**Figure S4.** Average leaf area curves of well-watered (solid lines) and 60% MWC water-stressed (dashed lines) birch (a), maple (b), and redbud (c) plants in experiment 1. From these curves, maximum leaf growth rate (d), day after treatment initiation on which leaves reached 50% of full expansion (e), and the number of days for leaves to reach full expansion from initiation (f). Data are means ± SE. * and ** denote a significant difference between WW and WS leaves at *P* < 0.05 and 0.01, respectively, based on one-way ANOVA (*n =* 4–6).

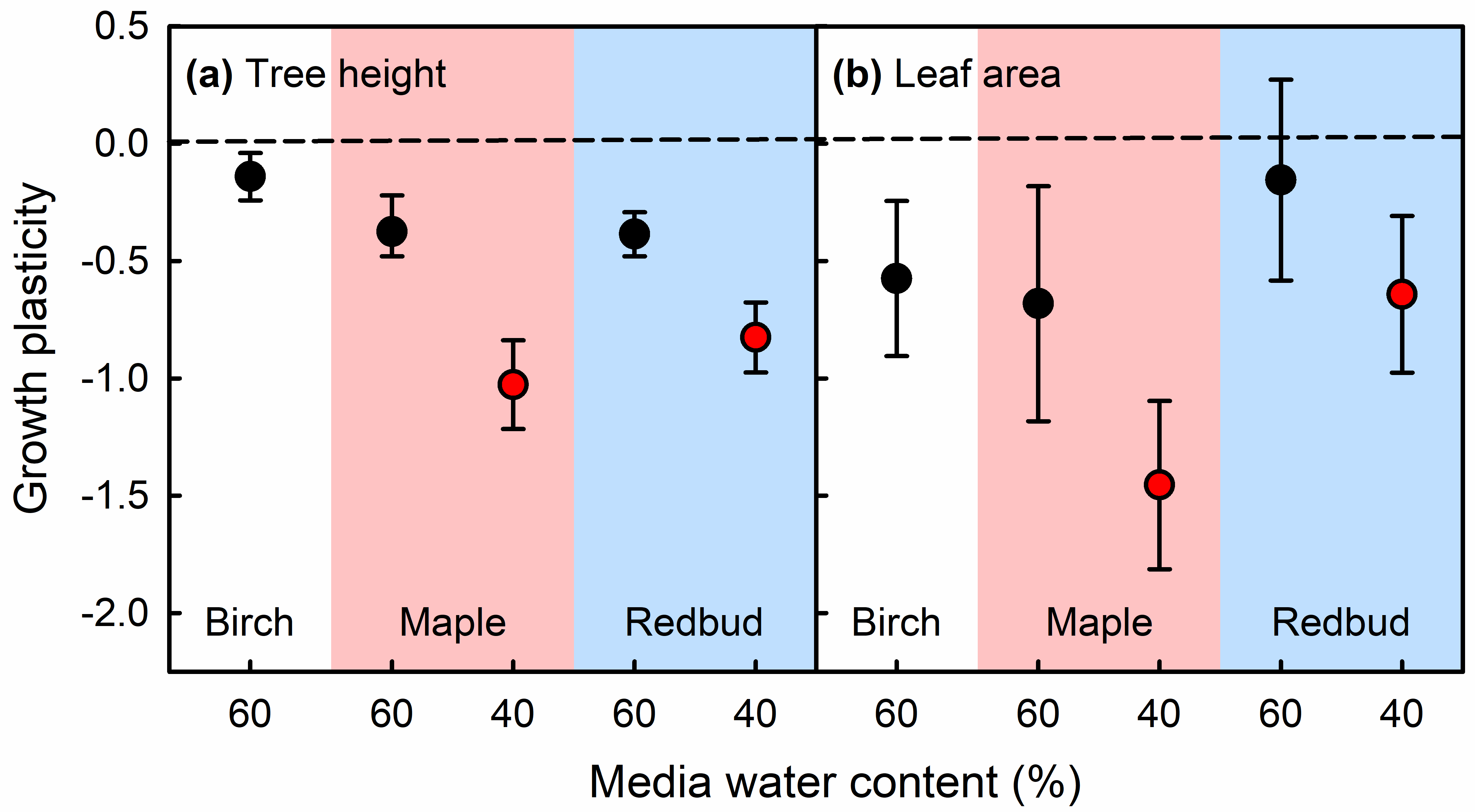

**Figure S5.** Plasticity of tree height (a) and individual leaf area (b) of plants that were grown under mild (60% MWC, black symbols) or severe (40% MWC, red symbols) water-stress (WS) treatments. Leaf area data were collected from three leaves that developed under the 60% MWC treatment or one leaf that developed under the 40% MWC treatment. Plasticity was calculated as the ln response ratio. Error bars are 95% confidence intervals. WS plants are significantly different from well-watered plants if bars do not overlap 0.

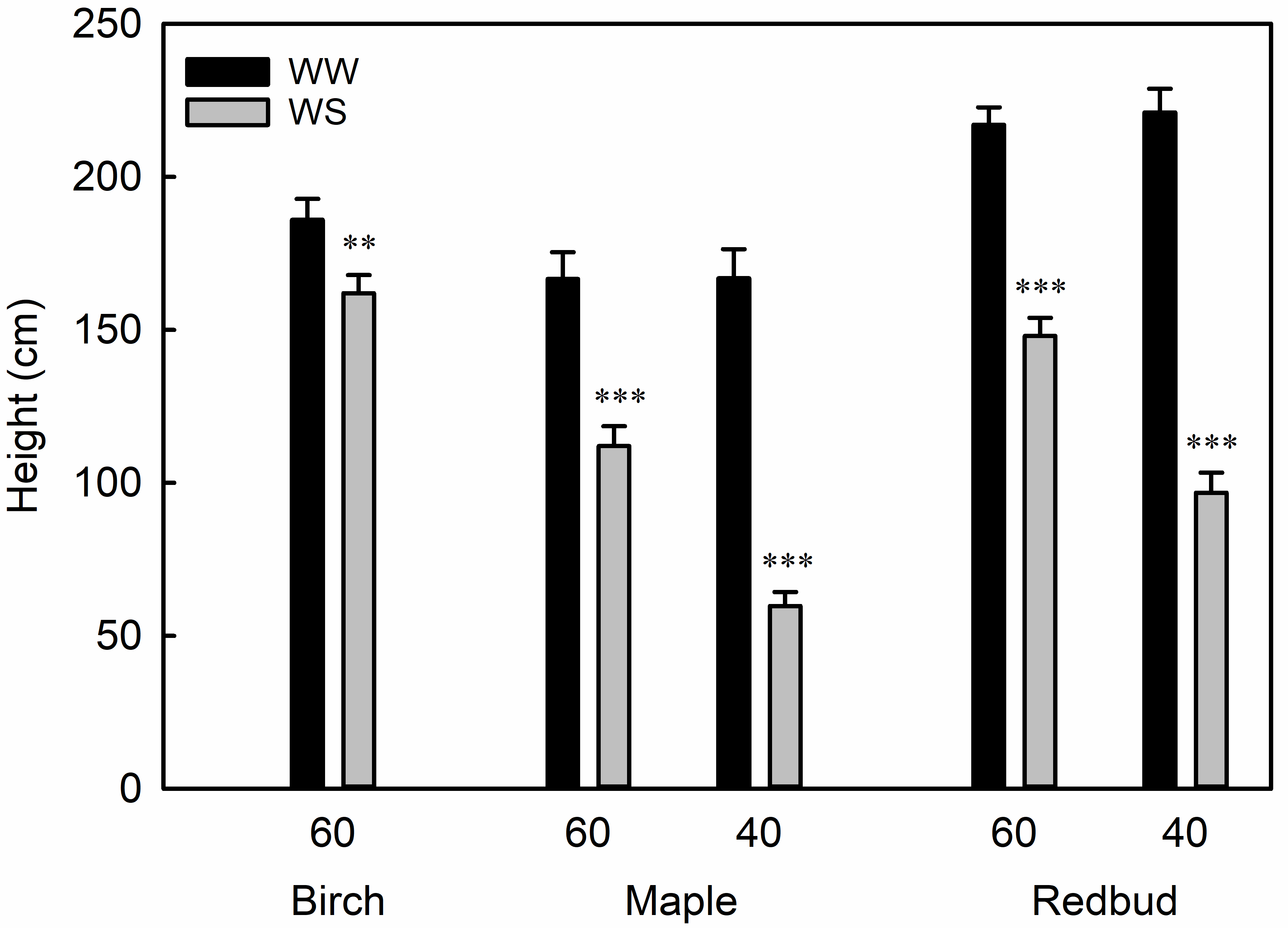

**Figure S6.** Height of trees after 82 (60% MWC) or 88 (40% MWC) days of growth under well-watered (WW) or water-stressed (WS, 60 or 40% MWC) conditions. Data are means ± SE, and ** or *** denote a significant difference between WW and WS trees within a species-treatment level at *P* < 0.01 or 0.001, respectively, based on one-way ANOVA (*n =* 4–6 for 60% MWC, *n =* 6–8 for 40% MWC).

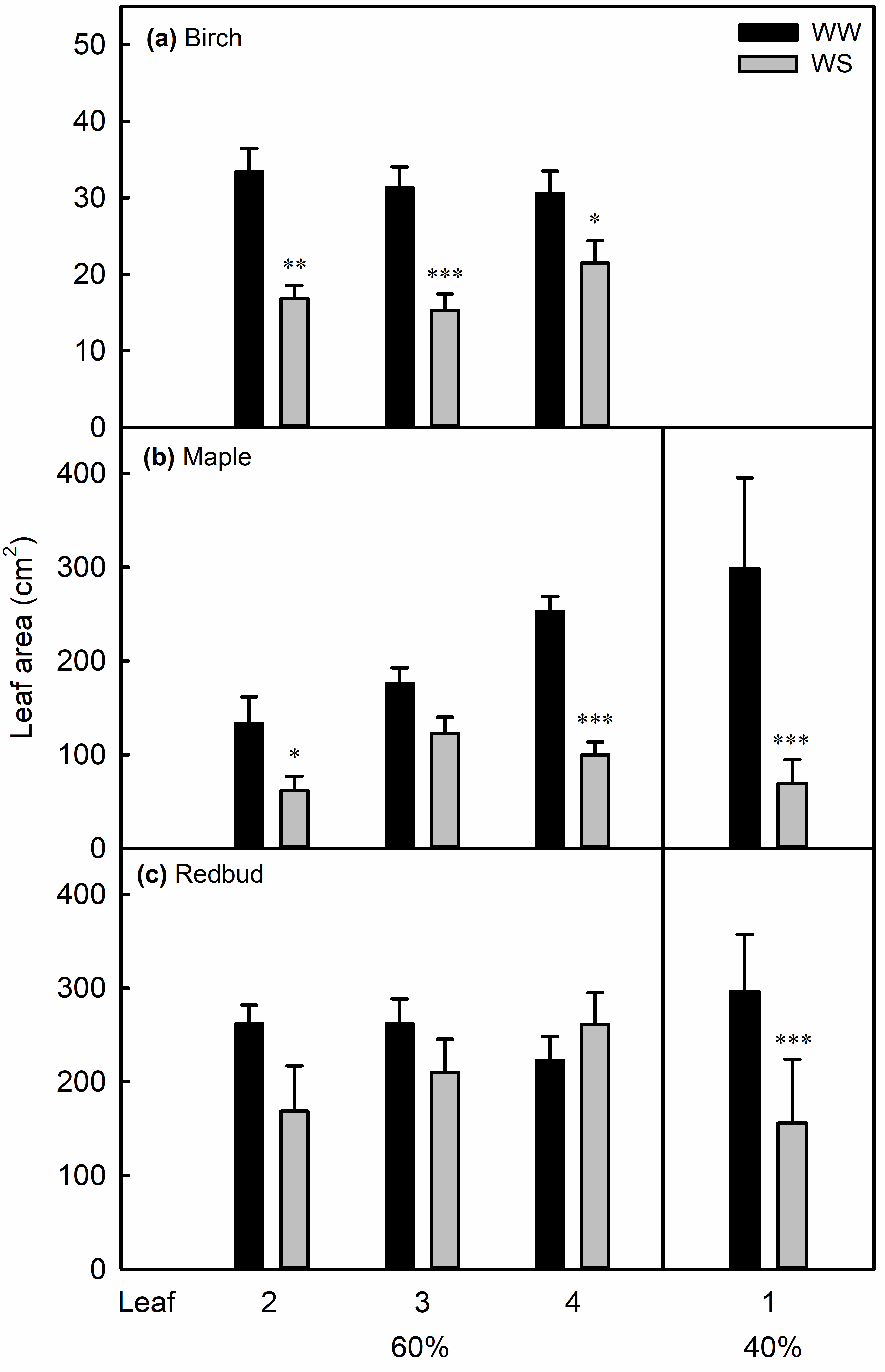

**Figure S7.** Leaf area of leaves 2, 3, and 4 or leaf 1 (see Fig. S3) measured after development under well-watered (WW) or water-stressed (WS, 60 or 40% MWC) conditions. Data are means ± SE, and *, **, or *** denote a significant difference between WW and WS leaves at *P* < 0.05, 0.01, or 0.001, respectively, based on one-way ANOVA (*n =* 4–6 for 60% MWC, *n =* 6–8 for 40% MWC).

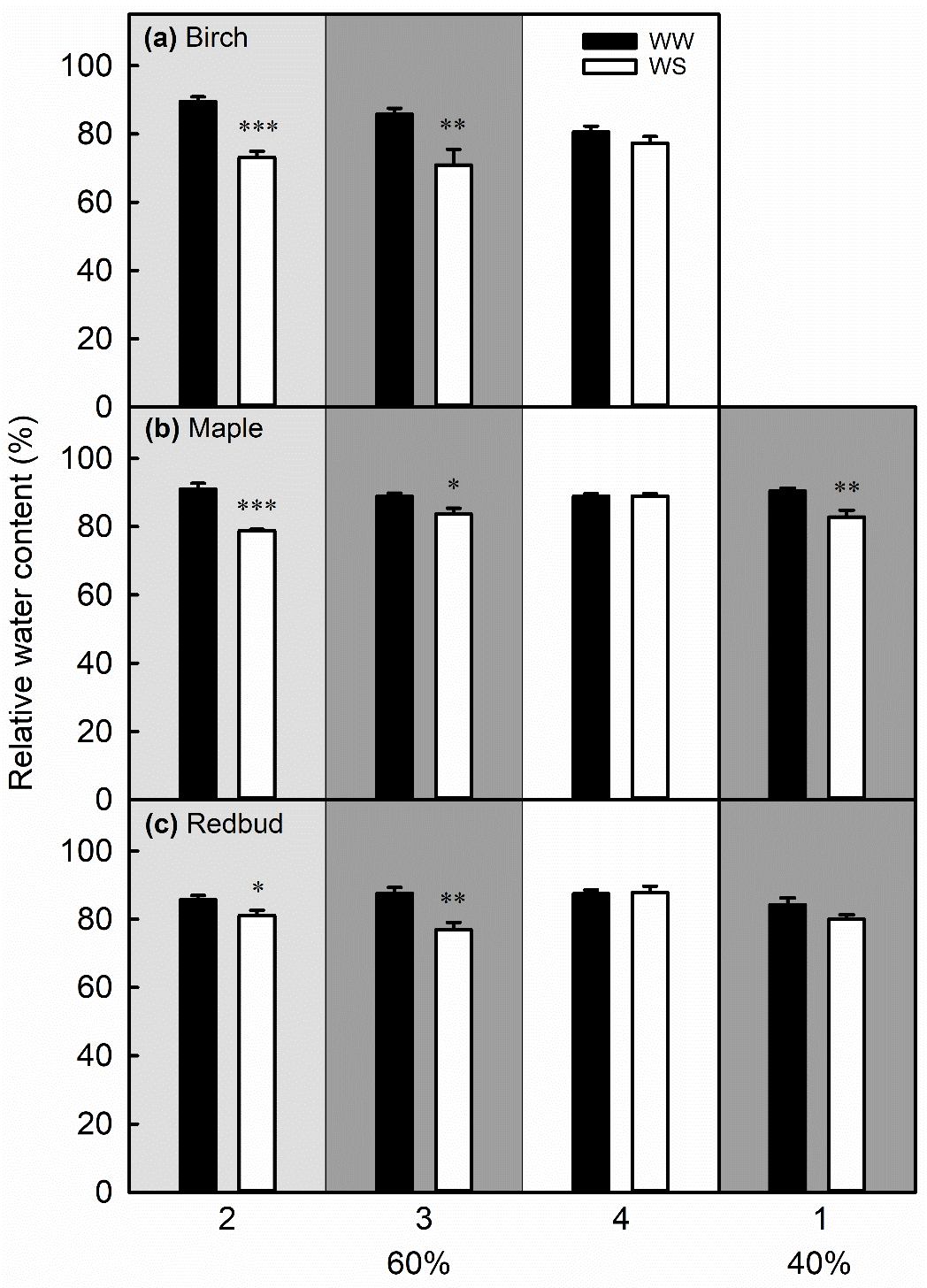

**Figure S8.** Relative water content of leaves 2, 3, and 4 or leaf 1 (see Fig. S3) that developed under well-watered (WW) or water-stressed (WS, 60 or 40% MWC) conditions. Background shading indicates the time that data were collected after the last irrigation: 18 h (light grey), 36 h (dark grey), or after re-irrigating (white). Data are means ± SE. In each time period, *, **, or *** denote a significant difference between WW and WS leaves at *P* < 0.05, 0.01, or 0.001, respectively, based on one-way ANOVA (*n =* 4–6 for 60% MWC, *n =* 6–8 for 40% MWC).

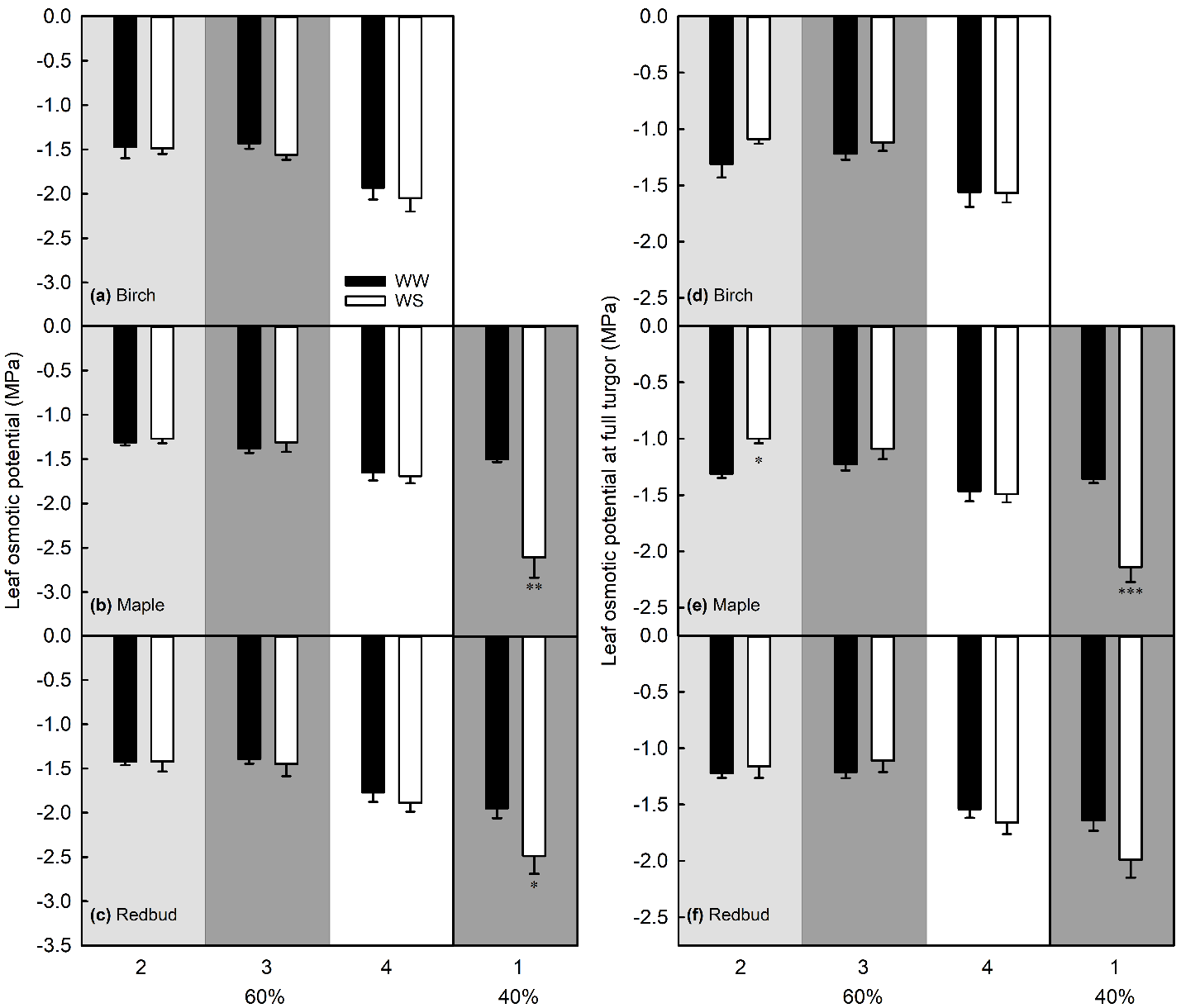

**Figure S9.** Leaf osmotic potential (Ψ_π_, a–c) and osmotic potential at full turgor (Ψ_π100_, d–f) of leaves 2, 3, and 4 or leaf 1 (see Fig. S3) that developed under well-watered (WW) or water-stressed (WS, 60 or 40% MWC) conditions. Background shading indicates the time that data were collected after the last irrigation: 18 h (light grey), 36 h (dark grey), or after re-irrigating (white). Data are means ± SE. In each time period, *, **, or *** denote a significant difference between WW and WS leaves at *P* < 0.05, 0.01, or 0.001, respectively, based on one-way ANOVA (*n =* 4–6 for 60% MWC and *n =* 6–8 for 40% MWC).

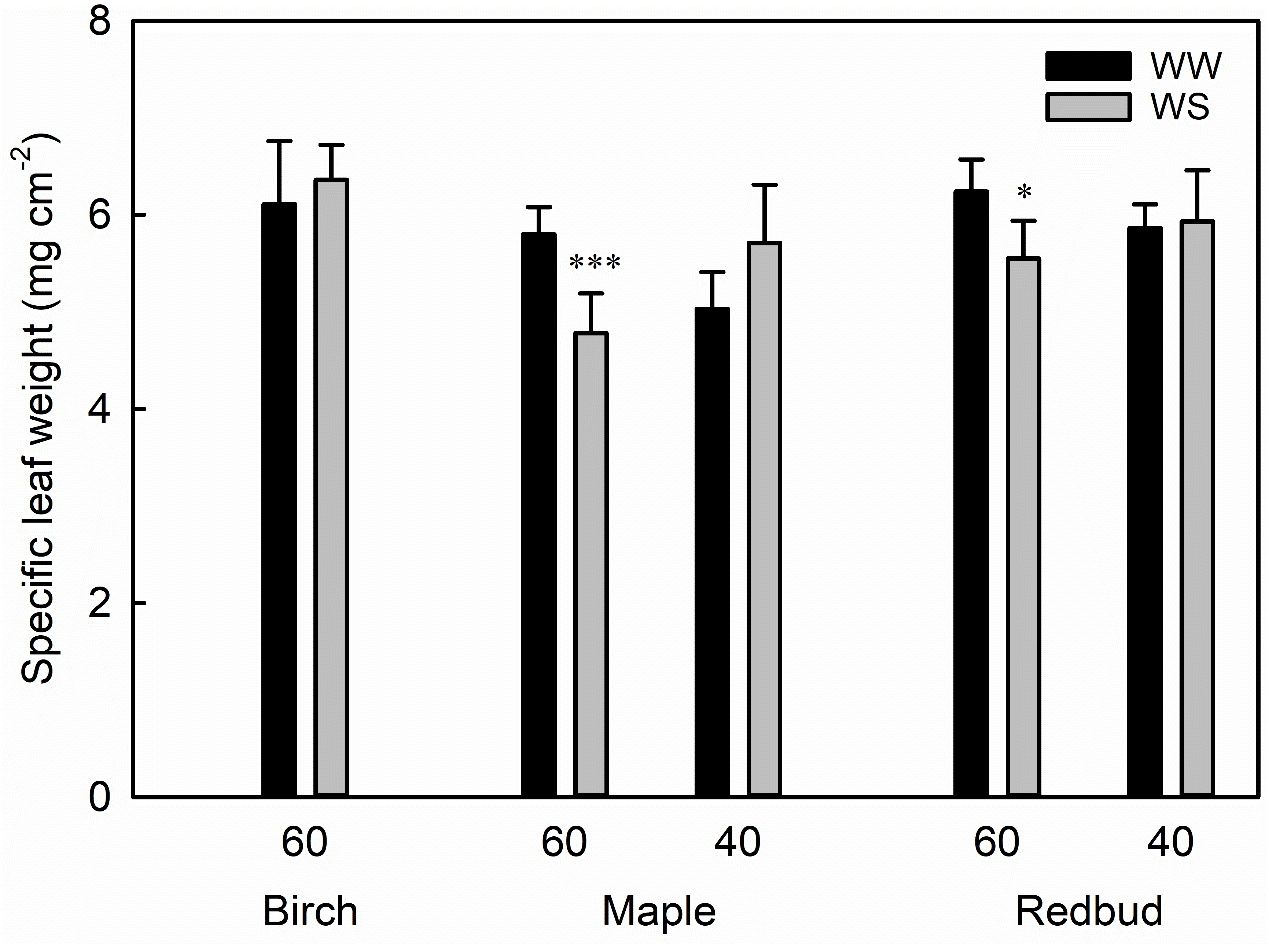

**Figure S10.** Specific leaf weight of leaves 2, 3, and 4 (see Fig. S3) measured after development under well-watered (WW) or water-stressed (WS, 60 or 40% MWC) conditions. Data are means ± SE, and * or *** denote a significant difference between WW and WS leaves at *P* < 0.05 or 0.001, respectively, based on one-way ANOVA (*n =* 4–6 for 60% MWC, *n =* 6–8 for 40% MWC).

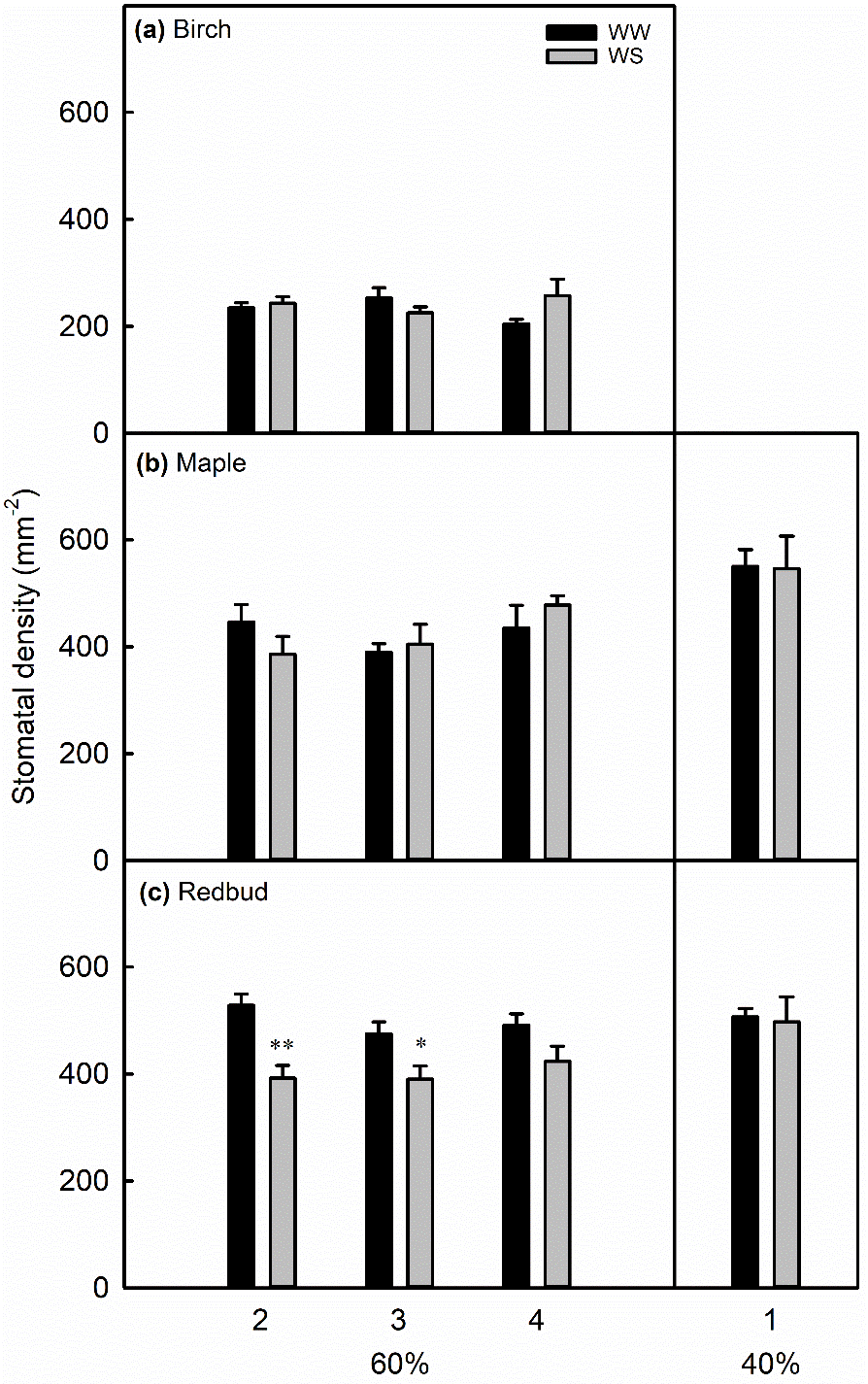

**Figure S11.** Stomatal density in leaves 2, 3, and 4 or leaf 1 (see Fig. S3) measured after development under well-watered (WW) or water-stressed (WS, 60 or 40% MWC) conditions. Data are means ± SE, and * or ** denote a significant difference between WW and WS leaves at *P* < 0.05 or 0.01, respectively, based on one-way ANOVA (*n =* 4–6 for 60% MWC, *n =* 6–8 for 40% MWC).

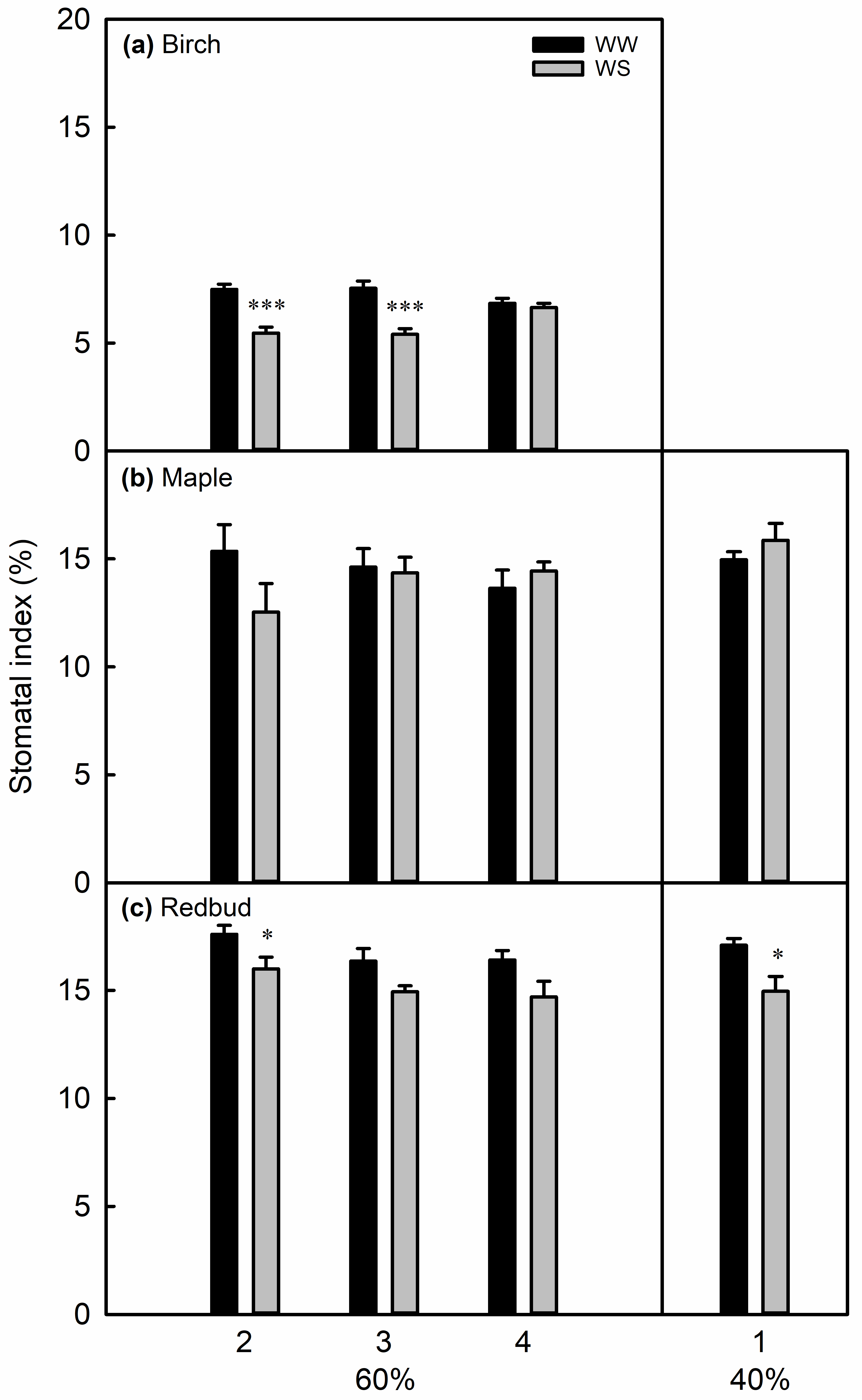

**Figure S12.** Stomatal index in leaves 2, 3, and 4 or leaf 1 (see Fig. S3) measured after development under well-watered (WW) or water-stressed (WS, 60 or 40% MWC) conditions. Data are means ± SE, and * or *** denote a significant difference between WW and WS leaves at *P* < 0.05 or 0.001, respectively, based on one-way ANOVA (*n =* 4–6 for 60% MWC, *n =* 6–8 for 40% MWC).

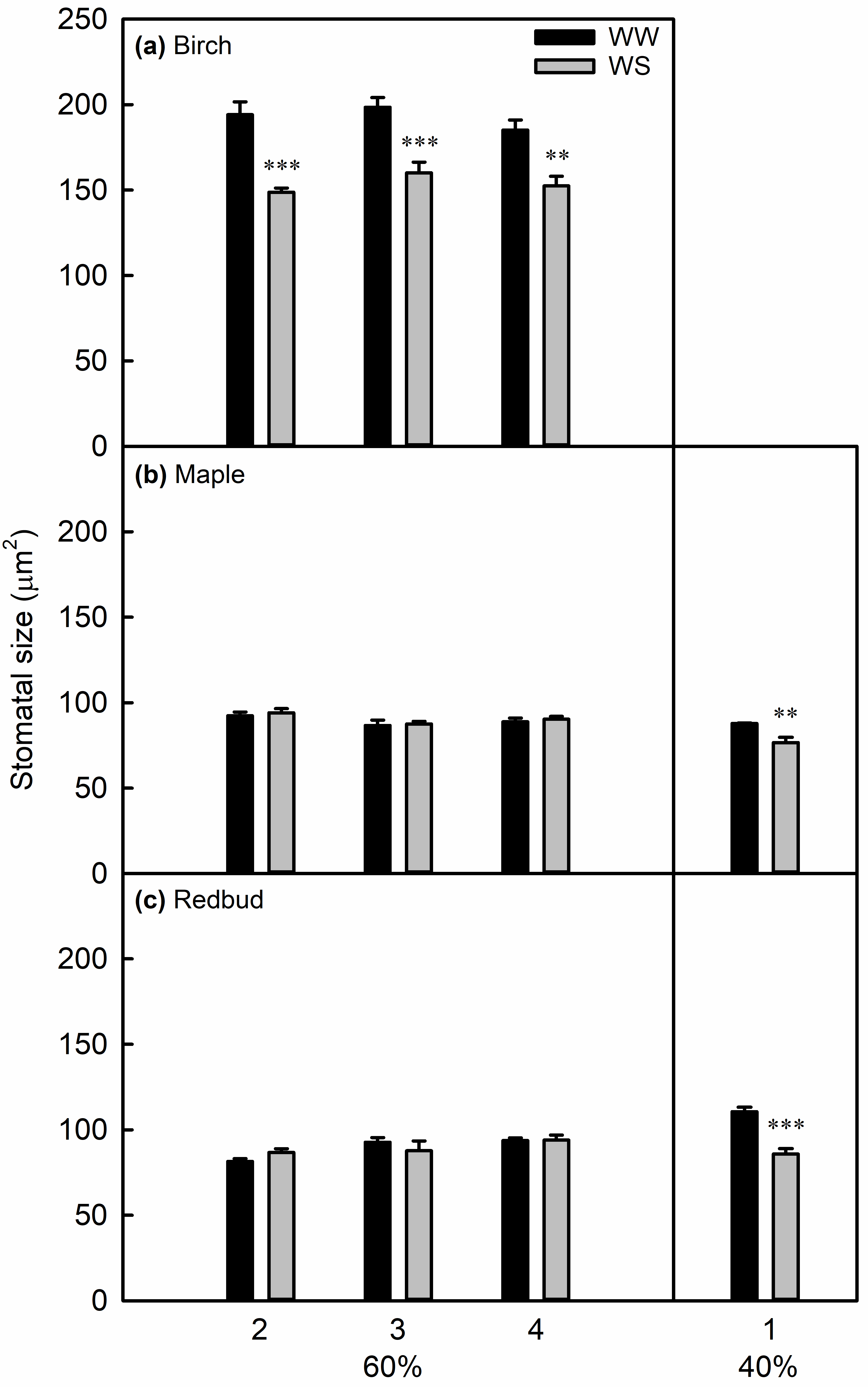

**Figure S13.** Stomatal size in leaves 2, 3, and 4 or leaf 1 (see Fig. S3) measured after development under well-watered (WW) or water-stressed (WS, 60 or 40% MWC) conditions. Data are means ± SE, and ** or *** denote a significant difference between WW and WS leaves at *P* < 0.01 or 0.001, respectively, based on one-way ANOVA (*n =* 4–6 for 60% MWC, *n =* 6–8 for 40% MWC).

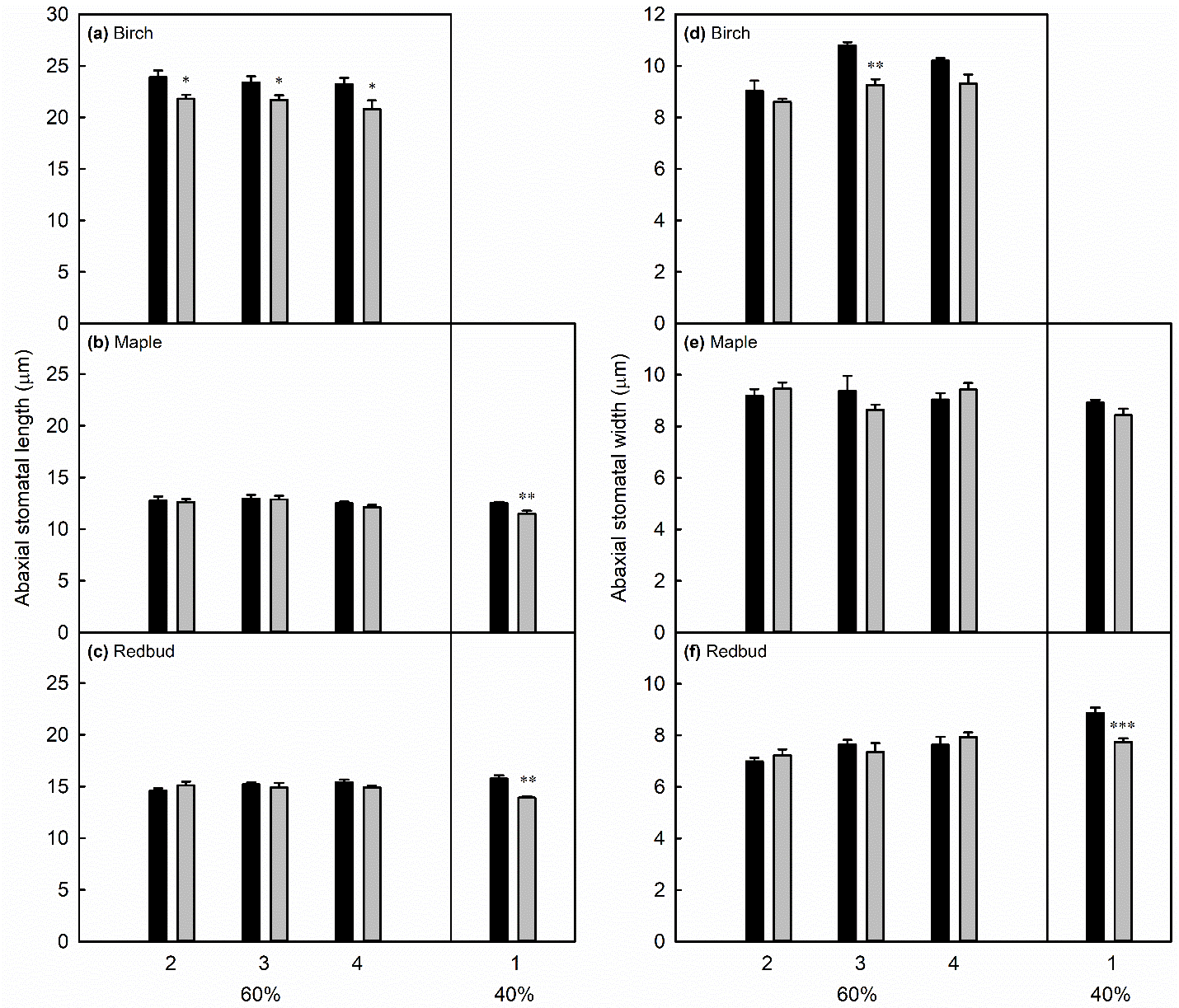

**Figure S14.** Stomatal length (a–c) and width (d–f) in leaves 2, 3, and 4 or leaf 1 (see Fig. S3) measured after development under well-watered (WW) or water-stressed (WS, 60 or 40% MWC) conditions. Data are means ± SE, and *, **, or *** denote a significant difference between WW and WS leaves at *P* < 0.05, 0.01, or 0.001, respectively, based on one-way ANOVA (*n =* 4–6 for 60% MWC, *n =* 6–8 for 40% MWC).

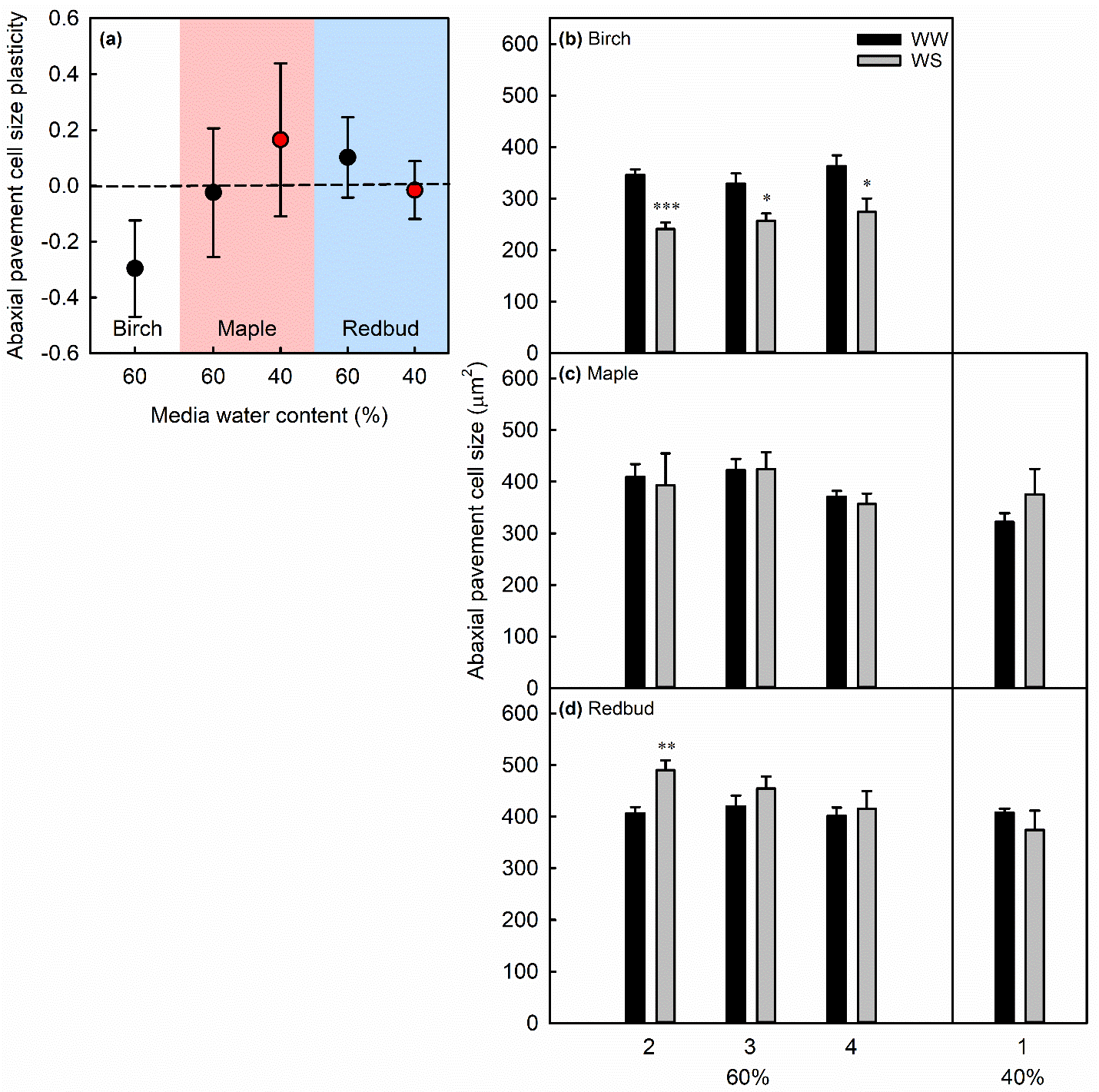

**Figure S15.** Plasticity of abaxial pavement cell size of leaves 2, 3, and 4 (see Fig. S3) that developed under mild (60% MWC, black symbols) or severe (40% MWC, red symbols) water-stress (WS) treatments (a). Plasticity was calculated as the ln response ratio. Data for individual leaves (b–d) are means ± SE, and *, **, or *** denote a significant difference between well-watered (WW) and WS leaves at *P* < 0.05, 0.01, or 0.001, respectively, based on one-way ANOVA (*n =* 4–6 for 60% MWC, *n =* 6–8 for 40% MWC).

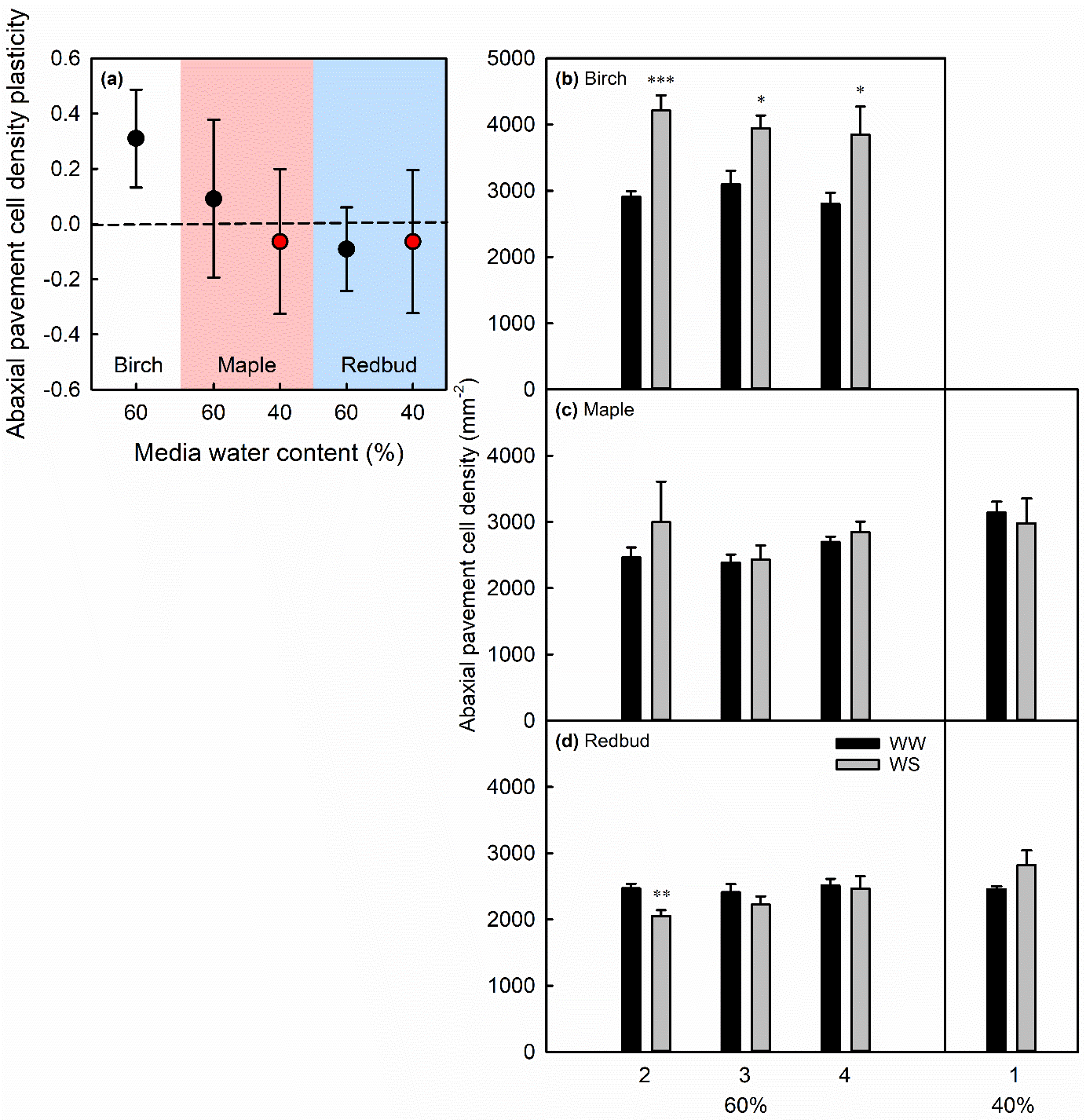

**Figure S16.** Plasticity of abaxial pavement cell density of leaves 2, 3, and 4 (see Fig. S3) that developed under mild (60% MWC, black symbols) or severe (40% MWC, red symbols) water-stress (WS) treatments (a). Plasticity was calculated as the ln response ratio. Data for individual leaves (b–d) are means ± SE, and *, **, or *** denote a significant difference between well-watered (WW) and WS leaves at *P* < 0.05, 0.01, or 0.001, respectively, based on one-way ANOVA (*n =* 4–6 for 60% MWC, *n =* 6–8 for 40% MWC).

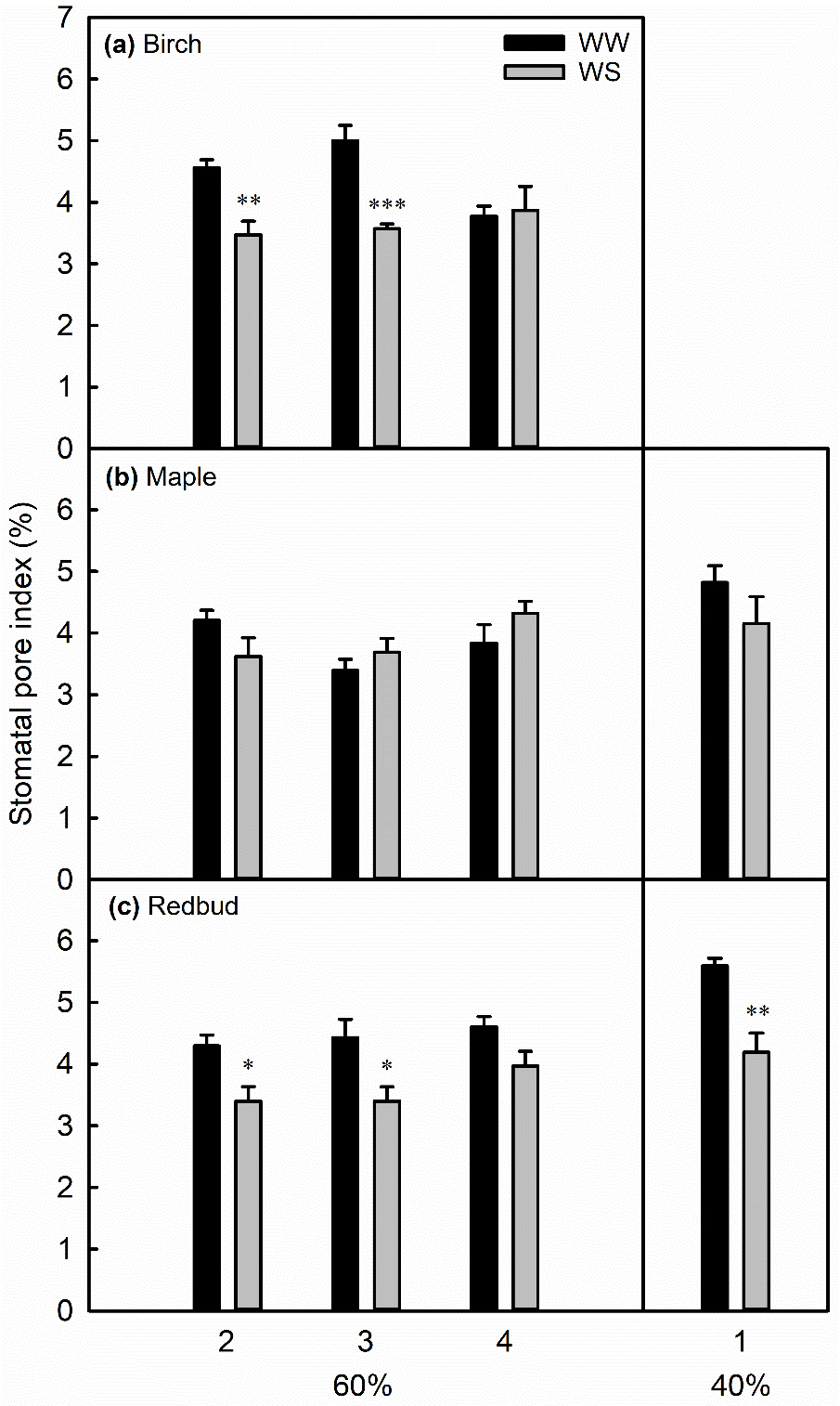

**Figure S17.** Stomatal pore index in leaves 2, 3, and 4 or leaf 1 (see Fig. S3) measured after development under well-watered (WW) or water-stressed (WS, 60 or 40% MWC) conditions. Data are means ± SE, and *, **, or *** denote a significant difference between WW and WS leaves at *P* < 0.05, 0.01, or 0.001, respectively, based on one-way ANOVA (*n =* 4–6 for 60% MWC, *n =* 6–8 for 40% MWC).

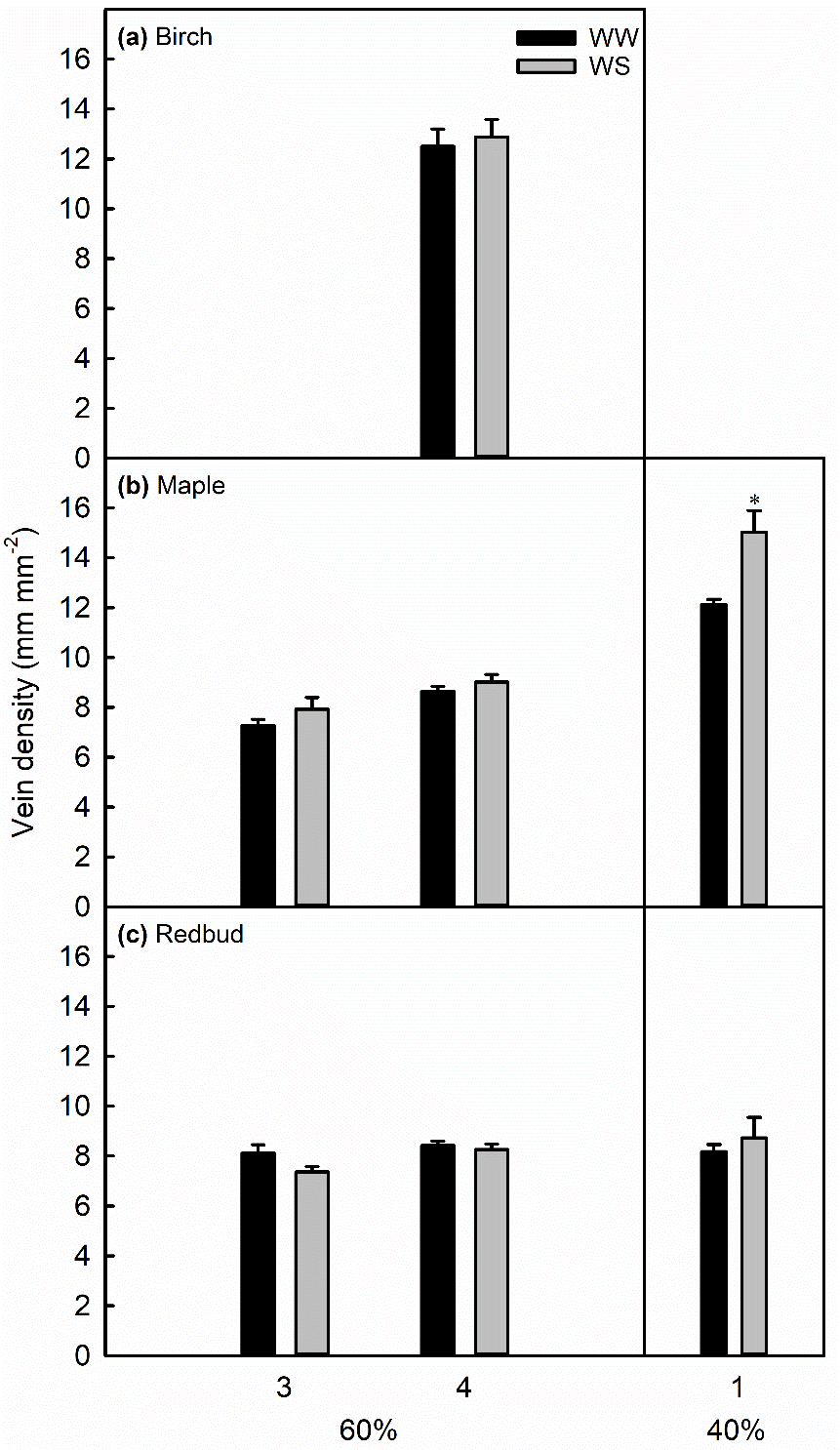

**Figure S18.** Vein density in leaves 3 and 4 or leaf 1 (see Fig. S3) measured after development under well-watered (WW) or water-stressed (WS, 60 or 40% MWC) conditions. Data are means ± SE, and * denotes a significant difference between WW and WS leaves at *P* < 0.05, based on one-way ANOVA (*n =* 4–6 for 60% MWC, *n =* 6–8 for 40% MWC).

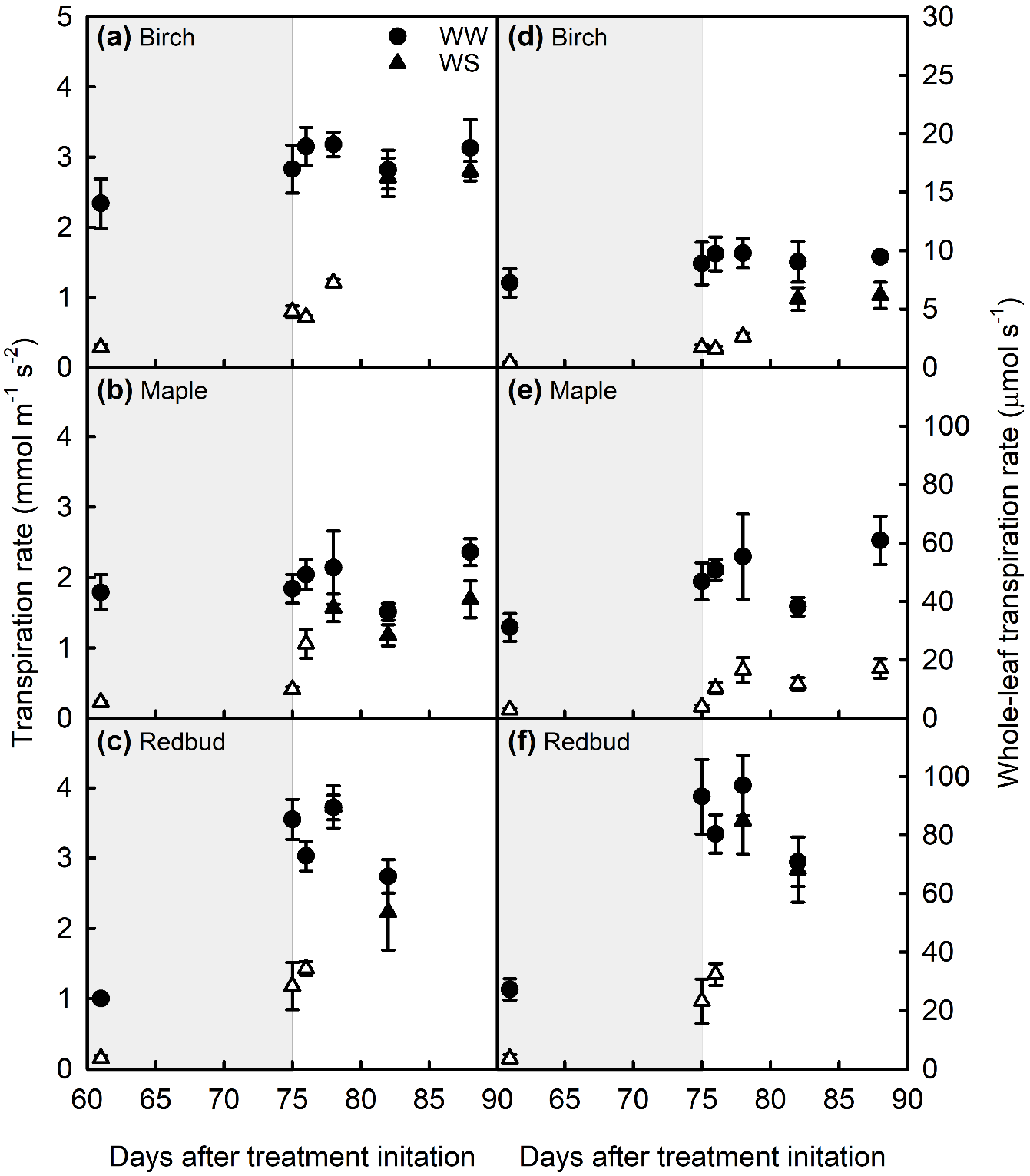

**Figure S19.** Transpiration rate (a–c) and whole-leaf transpiration rate (d–f) of leaves that developed under well-watered (WW) or water-stressed (WS, 60% MWC) conditions. Background shading indicates maintained WS over time (light grey) and re-irrigation (white). Leaf 3 was used for day 61 measurements and leaf 4 for all other measurements. Data are means ± SE. Open symbols indicate a significant difference between WS and WW plants at *P* < 0.05, based on one-way ANOVA (*n =* 4–6).

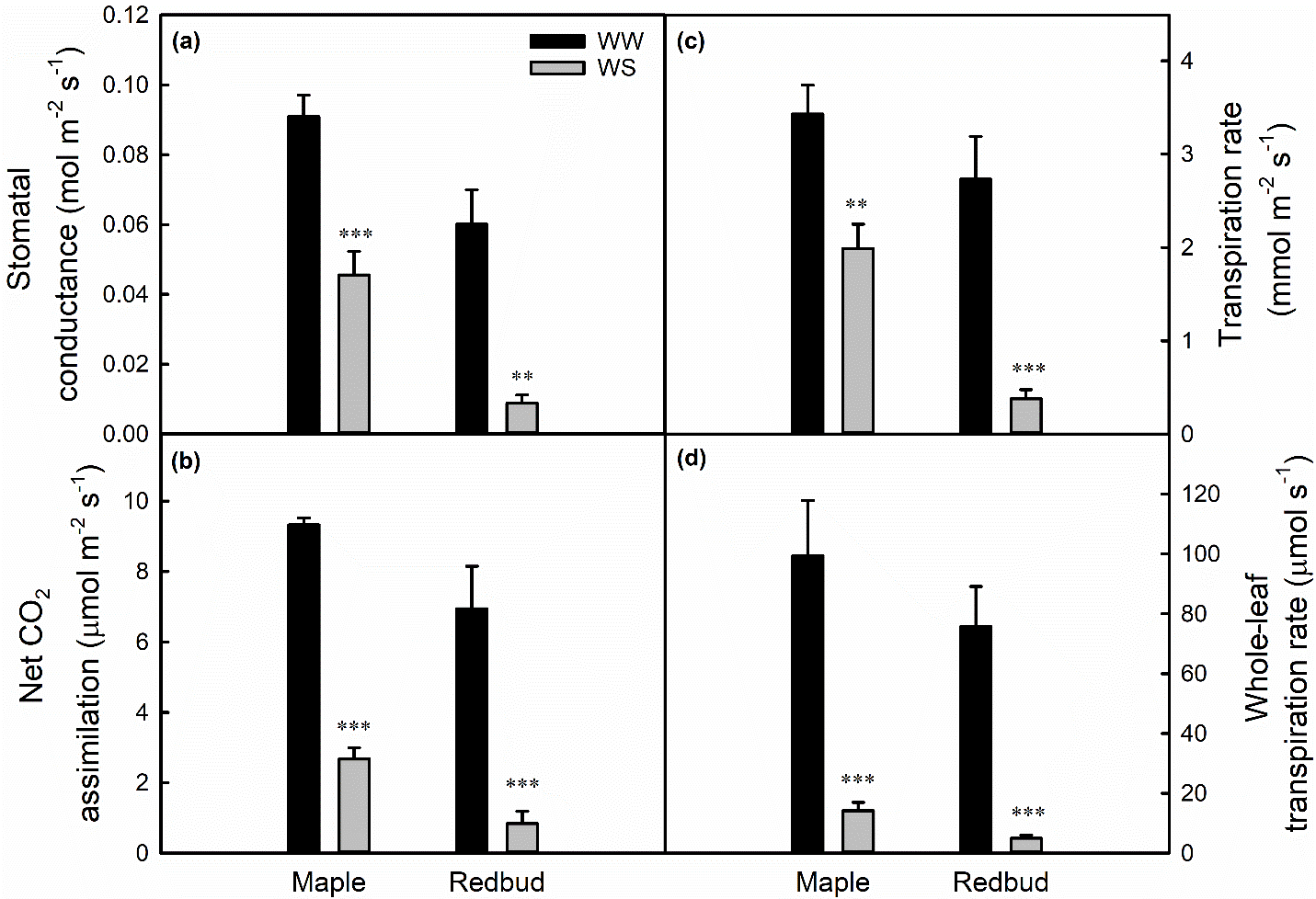

**Figure S20.** Stomatal conductance (a), net CO_2_ assimilation (b), transpiration rate (c), and whole-leaf transpiration rate (d), in maple and redbud leaves grown under well-watered (WW) or water-stressed (WS, 40% MWC) conditions. Data are means ± SE, and ** or *** denote a significant difference between WW and WS leaves at *P* < 0.01 or 0.001, respectively, based on one-way ANOVA (*n =* 6–8).

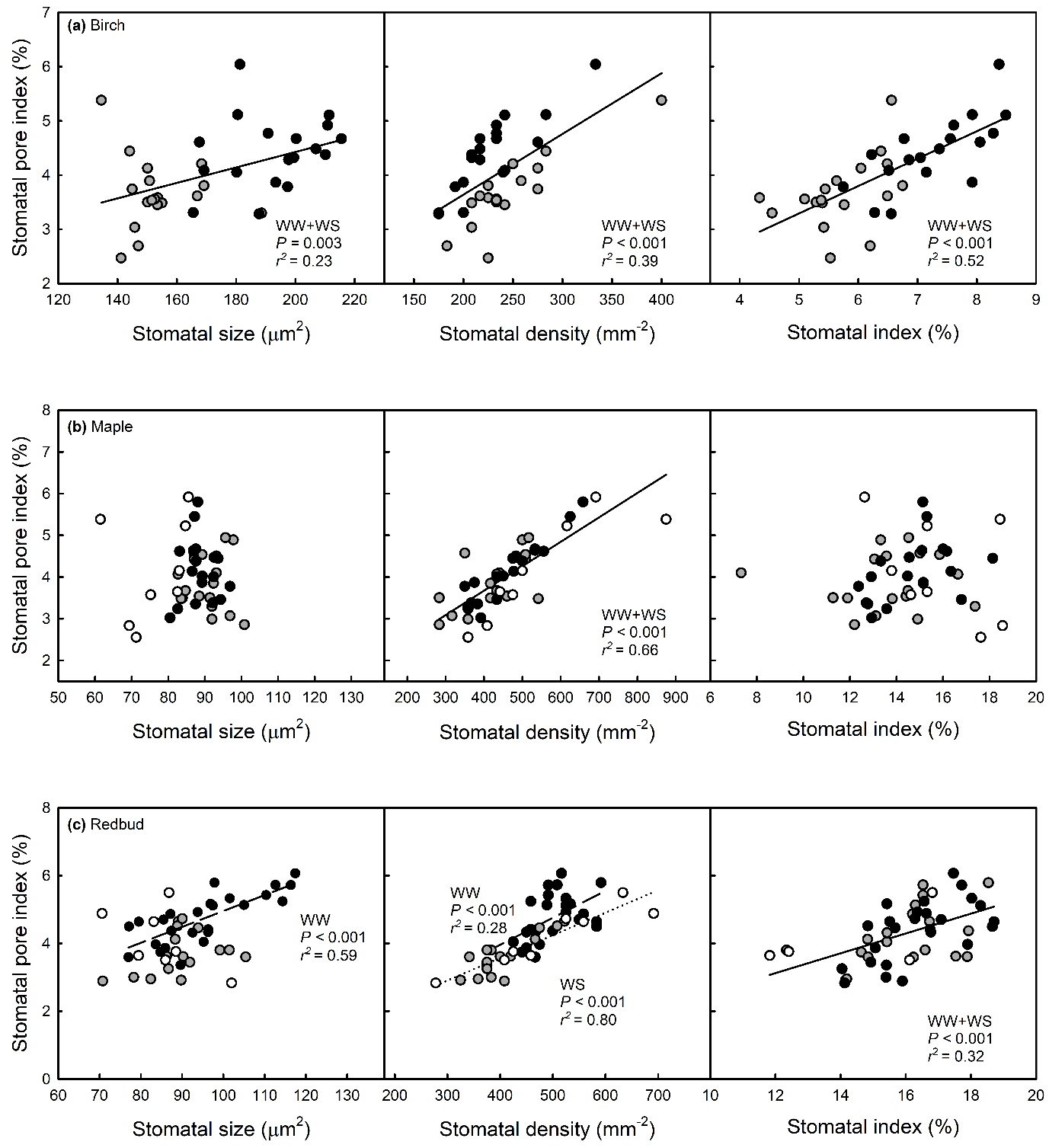

**Figure S21.** The relationship between stomatal pore index and stomatal size, density, and index. Regression lines, *P*-, and r^2^ values are reported for significant relationships at *P* < 0.05. In cases when treatments produced a significant difference in slope between well-watered (WW) and water-stressed (WS) treatments, regression lines are dashed for WW plants and dotted for WS plants. In cases when slopes were equal, solid black lines are used to show relationships between stomatal traits across all data points.

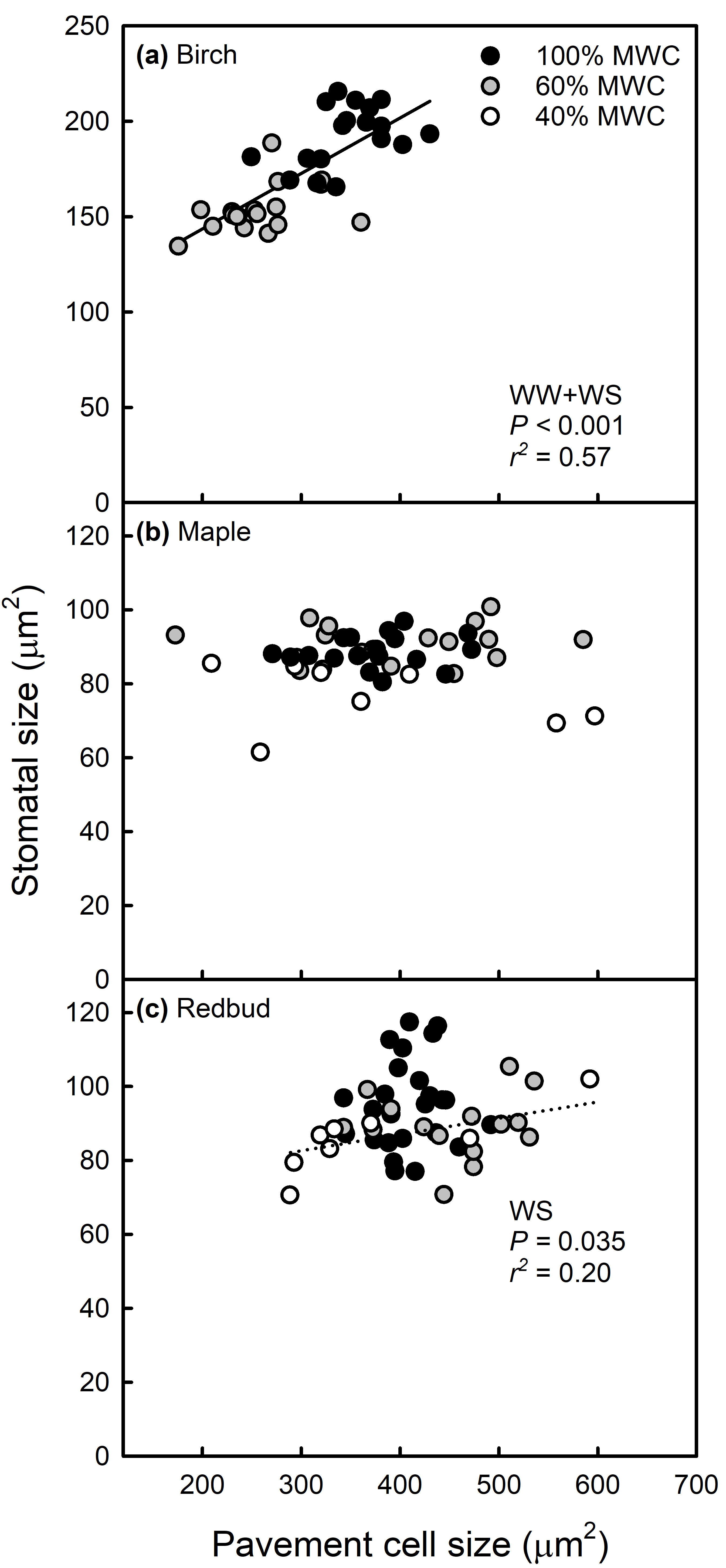

**Figure S22.** The relationship between pavement cell size and stomatal size. Regression lines, *P*-, and r^2^ values are reported for significant relationships at *P* < 0.05. Here, redbud and maple data are only shown for the 40% MWC treatment. In cases when treatments produced a significant difference in slope between well-watered (WW) and water-stressed (WS) treatments, regression lines are dashed for WW plants and dotted for WS plants. In cases when slopes were equal, solid black lines are used to show relationships between stomatal traits across all data points.

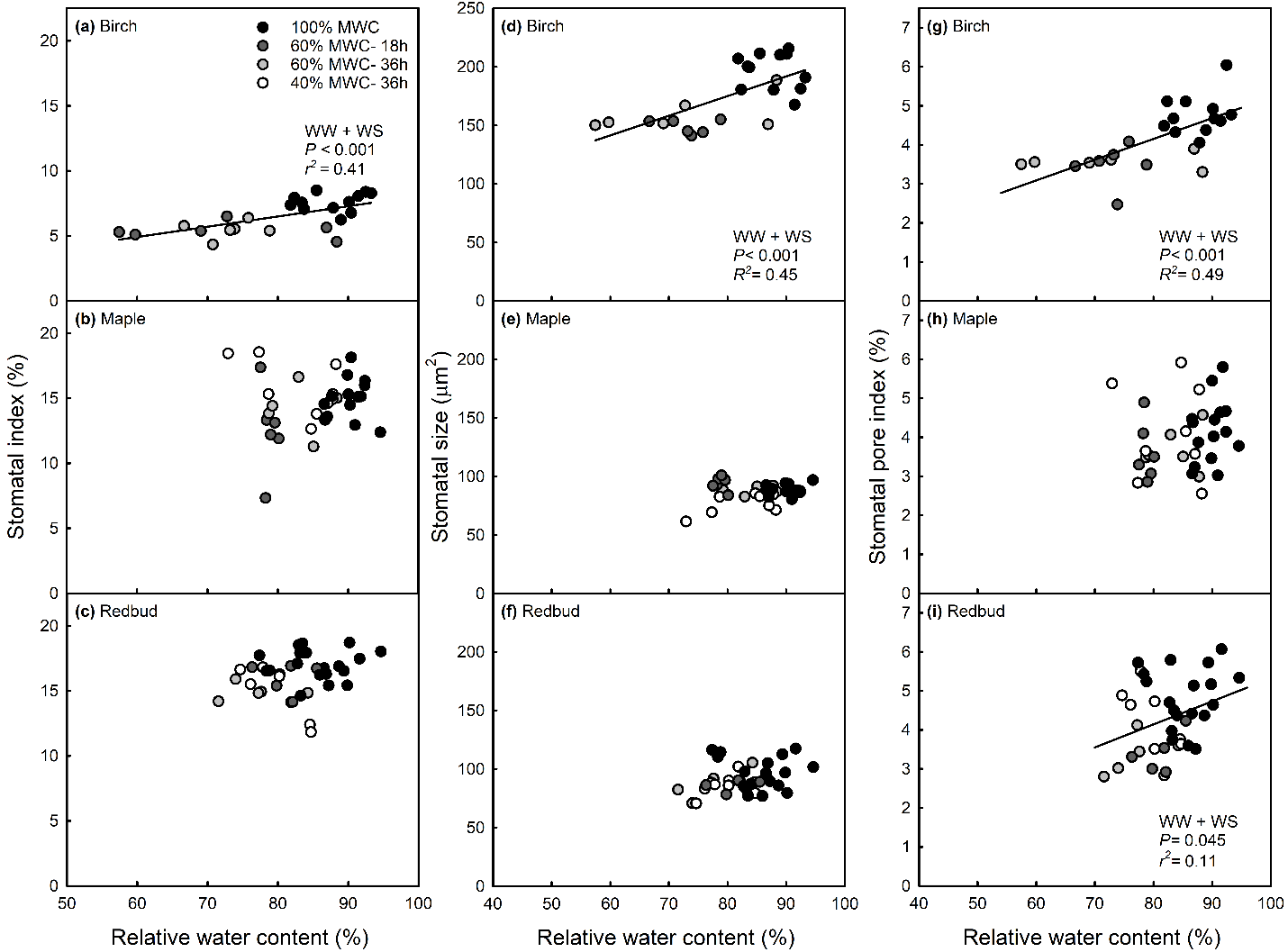

**Figure S2****3.** The relationship between stomatal index (a−c), size (d−f), and pore index (g−i) with relative water content (RWC) during the treatment period (i.e. pre-recovery). Media water content (MWC) and time since last irrigation are indicated in the legend. Regression lines, *P*-, and r^2^ values are reported for significant relationships at *P* < 0.05. Solid black lines are used to show relationships between water relations and stomatal traits across all data points.

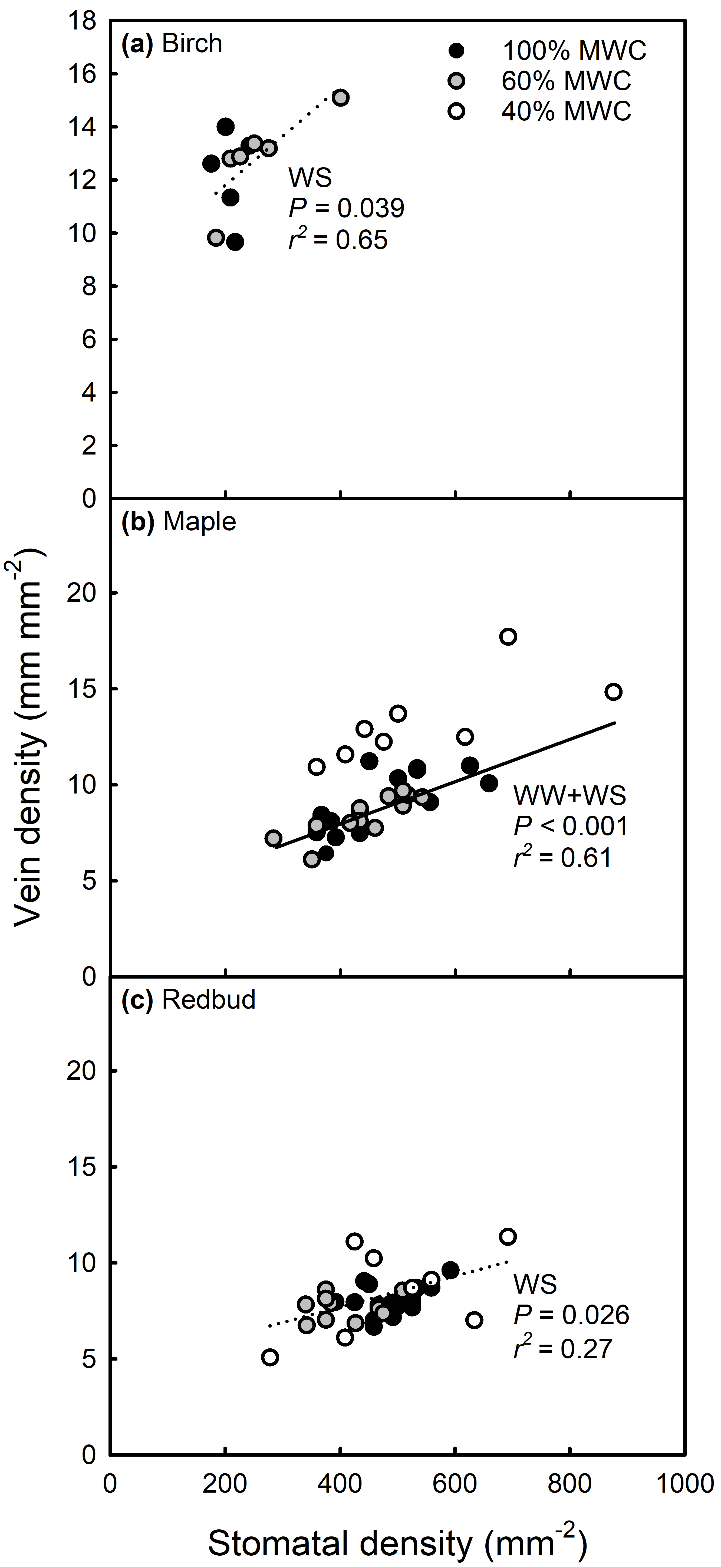

**Figure S24.** The relationship between stomatal and vein density. Regression lines, *P*-, and r^2^ values are reported for significant relationships at *P* < 0.05. In cases when treatments produced a significant difference in slope between well-watered (WW) and water-stressed (WS) treatments, regression lines are dashed for WW plants and dotted for WS plants. In cases when slopes were equal, solid black lines were used to show relationships between traits across all data points.

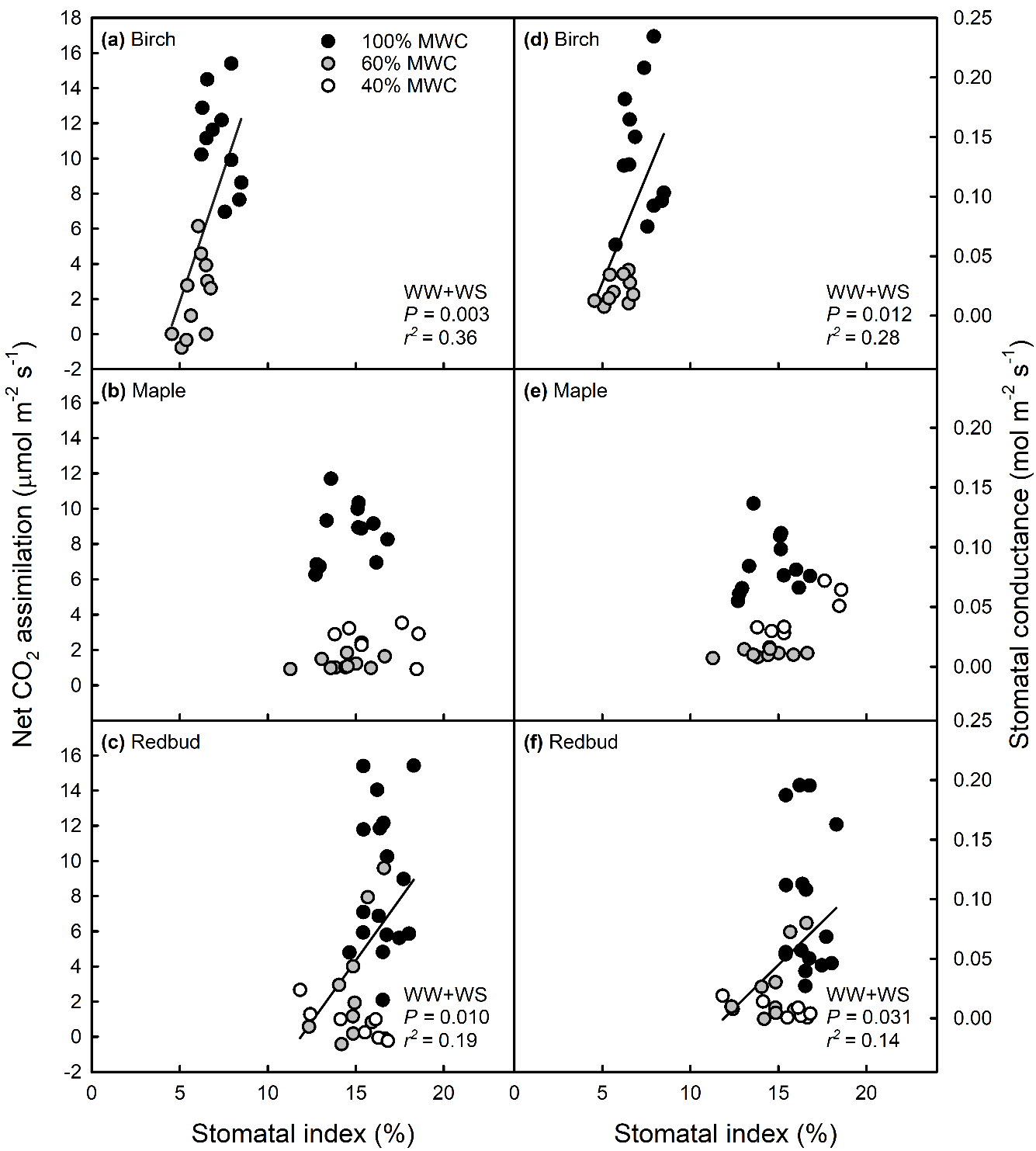

**Figure S25.** The relationship between stomatal index and net CO_2_ assimilation (left) and stomatal conductance (right). Gas exchange data are shown as collected during the stress period, excluding measurements made on leaves post-re-watering. Regression lines, *P*-, and r^2^ values are reported for significant relationships at *P* < 0.05. In cases when treatments produced a significant difference in slope between well-watered (WW) and water-stressed (WS) treatments, regression lines are dashed for WW plants and dotted for WS plants. In cases when slopes were equal, solid black lines are used to show relationships between stomatal traits across all data points.

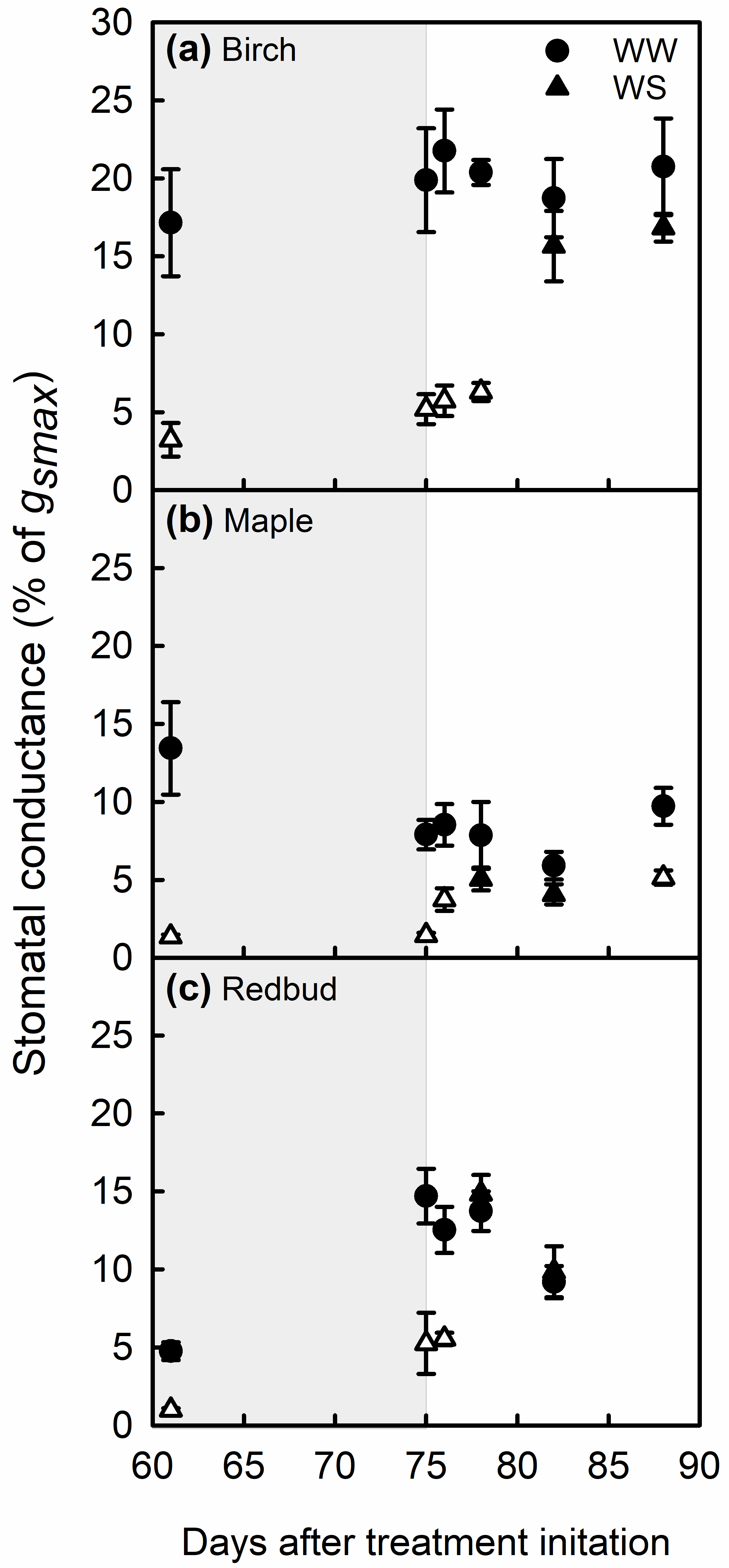

**Figure S26.** Changes in % of *g_smax_* in the fourth leaf that developed under well-watered (WW) or water-stressed (WS, 60% MWC) conditions. Shading within the plots indicates the treatment stage at the time of the measurement: maintained WS over time (light grey) and following recovery and daily irrigation to saturation (white). Data shown are means ± SE. White symbols indicate a significant difference between well-watered and (WS) plants at *P* < 0.05, based on one-way ANOVA (*n =* 4–6).

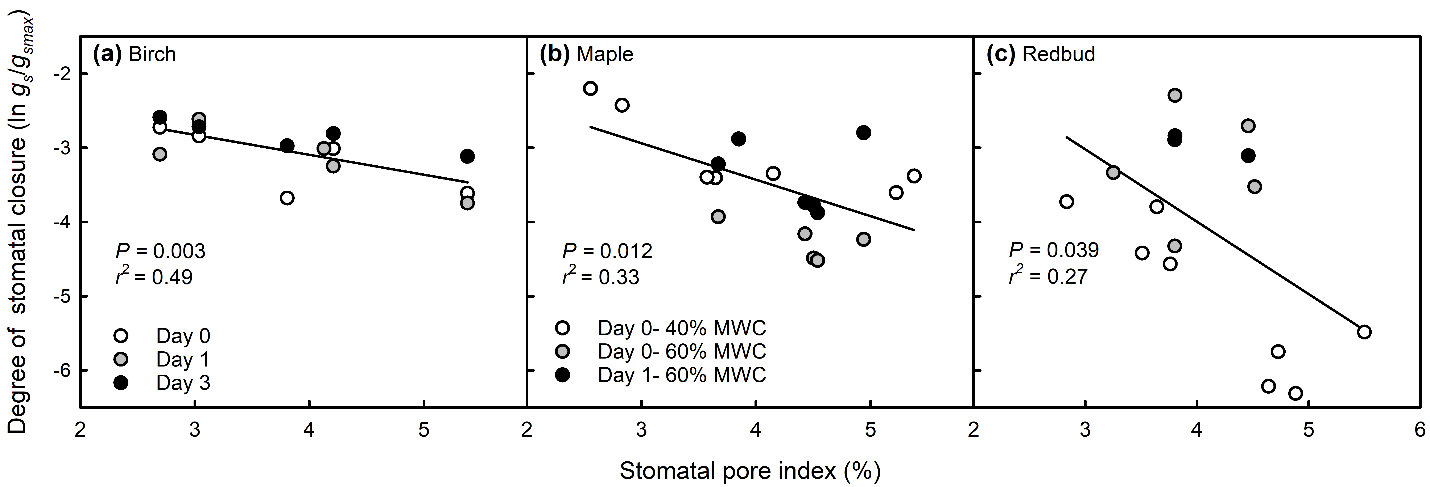

**Figure S27.** The relationship between the degree of stomatal closure (calculated as the ln of *g_s_* as a fraction of *g_smax_*) in WS plants and stomatal pore index. In birch leaves, data are shown for the fourth leaf that developed under the 60% MWC, for the day prior to re-watering as well as for two days after re-watering, prior to full recovery. In redbud and maple leaves, data are shown for the 40% MWC treatment and for the fourth leaf that developed under the 60% MWC, for the day prior and after re-watering. Regression lines, *P*-, and r^2^ values are reported for significant relationships at *P* < 0.05. Solid black lines are used to show relationships between water relations and stomatal traits across all data points.
